# Comparative Transcriptomics of Adherent and Suspension Chicken Fibroblast Cell Lines for the Optimization of Cultivated Meat Processes

**DOI:** 10.1101/2025.09.02.673824

**Authors:** Elizabeth J. Contreras, Archana Nagarajan, Benjamin H. Bromberg, Mason P. Villegas, David L. Kaplan

## Abstract

For the cultivated meat industry, food-relevant cell lines with optimal attributes for industrial bioprocesses are needed. One key trait is suspension-proficiency, or the ability of cells to proliferate in non-adherent culture conditions, to achieve high viable cell densities and streamline cell harvests. The limited success with suspension adaptations of animal cells for cultivated meat restricts the number of non-adherent cell lines suitable for cost-efficient biomass production. Here, we investigated transcriptional profiles of chicken embryonic fibroblasts (DF-1s) as they progressed from adherent to suspension culture using time-series RNA sequencing. From this analysis, we identified that DF-1s adapt to their non-adherent conditions by suppressing non-essential metabolism and tightly regulating cell junctions in response to oxidative stress and cell detachment. Based on these findings, we propose a novel mechanism enabling suspension proficiency in DF-1s, where mitogen-activated protein kinase (MAPK) p38δ sequesters TEA domain (TEAD) proteins to the cytoplasm and promotes Yes-associated protein (YAP) binding to Forkhead box protein 1 (FoxO1), simultaneously causing cell-matrix dissociation and antioxidant production. The outcome of this work is a deeper understanding of mechanisms that drive the generation of suspension cell lines, with a goal to translate these findings to other food-relevant cell lines and develop expanded cell options for the field.

## Introduction

Assuming that historical trends of food preferences continue, the demand for animal-sourced protein is expected to increase substantially across all World Health Organization regions over the next 25 years[1]. Poultry production is of notable significance due to the popularity of poultry-based foods; in 2018, 123 million metric tons of poultry meat were produced globally, with this number expected to rise by 15 million tons by 2027[2]. Cultivated meat, a branch of cellular agriculture, has the potential to diminish many of the negative environmental, animal welfare, and public health impacts associated with conventional meat production[3]. While many foundational studies have focused on the successful isolation of livestock cells, the adaptation of these cells to serum-free media, and the development of scaffolds to construct 3D cell-laden tissues, challenges concerning the scale-up of cultivated meat remain[4–10]. A key challenge is the lack of food-relevant, non-adherent cell lines suitable for the scale-up of cultivated meat.

Muscle, fat, and other food-relevant cells are inherently adherent due to the presence of integrins that bind cells to the extracellular matrix (ECM); these interactions are crucial for cell function within a living animal[11,12]. Integrins and other cell adhesion molecules (CAMs) are not only responsible for adhering cells to their substrate, but play a key role in differentiation as integrin expression changes during muscle and fat maturation[11,12].

However, non-adherent (i.e. suspension, anchorage-independent) cell lines are advantageous compared to adherent counterparts because they are capable of growing in industrial bioreactors, achieving higher viable cell densities (VCD), and simplifying the biomass harvest[13]. Suspension culture ranked highly in the most important cell line characteristics for cultivated meat production[13]. Despite advantages of anchorage-independent cells, very few suspension cell lines have been established for cultivated meat production. Difficulties in the generation of suspension cell lines stem from the lack of information about critical cellular and molecular changes involved in the transition between adherent and non-adherent growth states[14,15]. A lack of integrin binding to an extracellular matrix leads to changes in organization of the actin cytoskeleton and anoikis, a caspase-mediated apoptosis that cells undergo when detached or inadequately adhered to their substrate[15–17]. Factors like reactive oxygen species, kinase activation, hypoxia, and oncogene activation can give rise to anoikis-resistant cells which can survive in non-adherent conditions[16]. However, overcoming anoikis is not enough to drive cells into suspension; other factors such as changes in cell-matrix dissociation, cellular metabolism, and membrane protein expression also play a role in this transtion[15,18,19]. To obtain proliferative, suspension-proficient cells, all factors must be taken into consideration, making suspension adaptation a challenging and complex problem.

Generally, there are two methods to generate anchorage-independent cells: genetic engineering or non-targeted, “spontaneous” approaches that select for cells capable of proliferating in suspension. Thus far, only five food-relevant, proliferative cell lines have been adapted to suspension using spontaneous approaches, demonstrating the time-consuming, inefficient nature and high failure rate inherent to these methods. These cell lines include chicken embryonic fibroblasts, turkey satellite cells, bovine adipose-derived stem cells, porcine fibroblasts, and porcine myoblasts [4,20–22]. Of these, spontaneously-adapted suspension turkey satellite cells and bovine adipose-derived stem cells survive, but do not proliferate, in non-adherent conditions, negating the advantages of suspension culture[21,22]. For the other cell types, mechanisms of suspension adaptation remain unclear, as they were not explored in previous studies.

Additionally, the non-targeted nature of these approaches further obscures the translatability of these findings. Alternatively, adherent cells may be adapted to suspension via genetic engineering, which was achieved in two bovine cell lines by knocking down genes that would confer anoikis resistance, such as *PTEN*[23,24]. Though the expression of *PTEN*, caspase genes such as *CASP3*, and integrin genes such as *ITGB1* were explicitly reduced in these engineered cell lines, other genes and pathways may also contribute to the ability of the cells to proliferate in suspension culture[23]. For example, there may be cellular changes to compensate for the loss of gene expression or downstream effects beyond the initial gene knockouts; this effect has been shown by transcriptome-wide changes following a single gene knockout[25]. Thus, even in genetically engineered suspension cell lines, there is a lack of foundational knowledge into mechanisms which contribute to proliferation of mammalian cells in anchorage-independent conditions.

Multiple publications have performed comparative transcriptomics between anchorage-dependent and anchorage-independent cells; however, none have focused on food-relevant cell lines. For example, suspension-adapted baby hamster kidney (BHK), Chinese hamster ovary (CHO), and Vero cells have been compared to their adherent counterparts utilizing RNA sequencing[17,19,26–28]. These studies found a plethora of genes which were downregulated in suspension cell lines, some of which were expected and somewhat ubiquitous, such as integrins (e.g. *ITGB3*), and others which were not, such as aquaporins (e.g. *AQP1*)[17,28]. Other studies have instead focused on performing analysis of larger datasets which include many adherent and suspension cell lines. For example, transcriptomic changes were evaluated between 141 anchorage-dependent and 39 anchorage-independent human cell lines[18,29]. Similarly, parental HEK293 cell lines were compared to anchorage-independent progeny lines, finding changes in gene expression that were also observed in 47 adherent and 16 suspension human cell lines[14]. Though these studies are foundational in discovering translatability across human cell lines, the non-adherent cell lines used to validate this pattern were human derived. Thus, relevance to animal cell lines used for cultivated meat production remains to be determined[14].

To address the gap, our lab adapted a commercially available chicken (*Gallus gallus*) embryonic fibroblast cell line, termed DF-1, to suspension culture using a published adaptation method[4,30]. While this prior published work included RNA sequencing (RNA-seq) of chicken fibroblast lines, the analysis compared primary, adherent cells to immortalized, suspension cells, confounding whether gene and pathway enrichment was driven by anchorage-independence or immortalization. Additionally, the analysis only included the temporal extremes of the adaptation process, which could further obscure transitional dynamics or nonlinear gene expression over time. In the present study, we utilized DF-1s as a model cell line to study suspension adaptation in order to provide insight into the differences between adherent and non-adherent cell lines at the mechanistic level. Using differential gene expression analysis (DGEA), gene set enrichment analysis (GSEA), and Mfuzz clustering techniques, we found that DF-1 cells early in the adaptation process responded to oxidative stress and loss of cell-matrix interactions through the transient activation of focal adhesions and pro-angiogenic signaling while simultaneously suppressing DNA replication. However, sustained downregulation of non-essential lipid metabolism and cell adhesion molecules allow energy reallocation toward increased cellular communication, representing the long-term adaptation strategy of suspension DF-1 cells. Non-adherent DF-1s at the end of the pipeline showed a sustained transcriptional profile that was similar to the adherent baseline, indicating recovery, resilience, and stability.

## Results

### 1. Non-Adherent DF-1 Cells Acquire Distinct Cellular Properties and Scale-Up Potential

Anchorage-dependent DF-1 cells were adapted to suspension culture by following a systematic pipeline[4]. This established methodology consists of five granular steps, allowing cells to gradually transition to suspension using selection pressure which enriches for anoikis-resistant, non-adherent, proliferative single cells (Fig. 1A).

**Figure 1:**
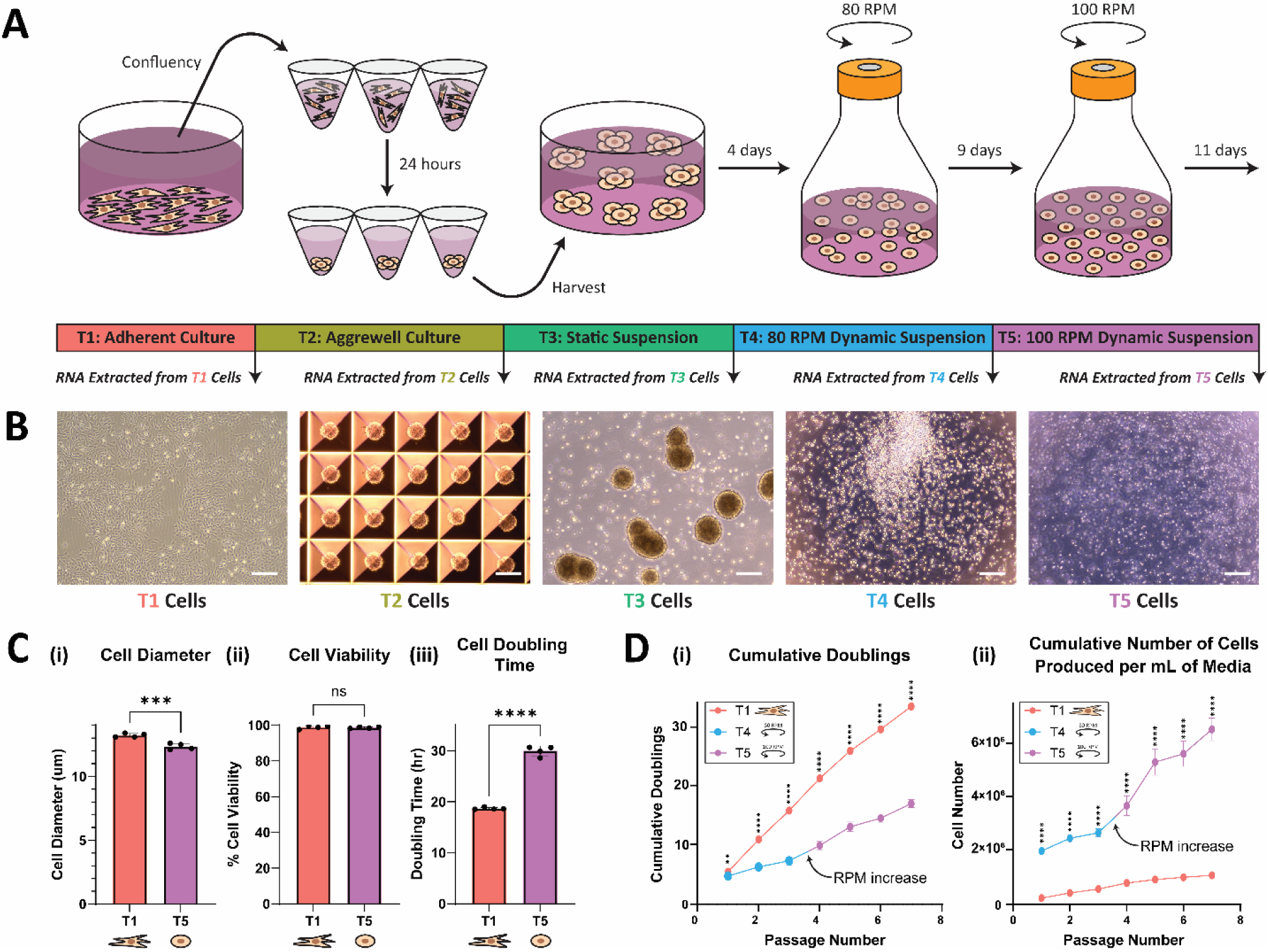
Non-adherent DF-1 cells acquire distinct cellular properties and scale-up potential. (A) Schematic diagram of suspension adaptation method adapted from Pasitka et al[4]. (B) Phase contrast images of DF-1 cells at each distinct time point during suspension adaptation. Scale bars for all images equal 240 μm. (C) Comparison between (i) cell diameter, (ii) cell viability, (iii) and cell doubling time of DF-1s before and after undergoing suspension adaptation. Error bars represent standard deviation, n = 4; statistical significance calculated by unpaired t-test, in which ns indicates no statistical significance, *** indicates p < 0.001, and **** indicates p < 0.0001. (D, i) Cumulative cell doublings of adherent and suspension DF-1s over 7 passages. Error bars represent standard deviation, n = 3 or 4; statistical significance calculated by multiple unpaired t-tests, in which ** indicates p < 0.01, and **** indicates p < 0.0001. (D, ii) Cumulative number of adherent or suspension DF-1s produced per mL of media over 7 passages. Error bars represent standard deviation, n = 3 or 4; statistical significance calculated by multiple unpaired t-tests, in which **** indicates p < 0.0001.

#### 1.1 Suspension Proficiency Leads to Distinct Morphological and Growth Characteristics

Over the course of the adaptation process, distinct morphological changes were observed at each of the five time points: adherent culture (T1), spheroid culture (T2), static suspension culture (T3), dynamic suspension culture at 80 revolutions per minute (RPM) (T4), and dynamic suspension culture at 100 RPM (T5) (Fig. 1B). At the first stage in the protocol, DF-1s displayed characteristic fibroblastic morphology (i.e. elongated and spindle-shaped cells) with a diameter averaging 13.2 μm and a doubling time of 18.65 hours (Fig. 1Ci, iii). Adherent “T1” cells were then seeded to form uniform, anchorage-independent “T2” spheroids averaging around 200 μm in diameter (Fig. 1B). These spheroids were cultured in static suspension and mechanically disturbed via daily micro-pipetting. Despite mechanical dissociation, DF-1 spheroids aggregated into large, irregular clusters of “T3” cells over time (Supplementary Fig. 1A). These large, non-uniform aggregates were then cultured in shaker flasks at 80 RPM; this dynamic culture condition causing “T4” cell clusters to break apart. Within shaker flasks, single cells as well as aggregates were observed (Supplementary Fig. 1B). To obtain a single-cell suspension, the shaking speed was increased to 100 RPM to break up the remaining aggregates via shear stress[31]. At this point, the “T5” cells were considered suspension-adapted due to their consistent average doubling time of 29.93 hours with a viability comparable to the initial adherent DF-1s (Fig. 1Cii, iii). Notably, the cell diameter of these suspension cells was significantly decreased to an average of 12.4 μm compared to their adherent counterparts, perhaps reflecting changes in metabolism or limitations due to loss of adherence (Fig. 1Ci).

#### 1.2 Altered Growth Characteristics Enable Bioprocess Intensification Through VCD

Anchorage-independent DF-1s exhibited slower doubling times than adherent DF-1s, resulting in reduced cumulative doublings over the same time period (Fig. 1Di). However, the relative viable cell yield of suspension DF-1s, measured as the cumulative number of cells produced per mL of media, remained significantly higher (Fig. 1Dii). This counterintuitive result is due to the higher maximum VCD, which is only possible in suspension conditions where cell growth is limited by volume rather than surface area. Because cell culture media is a cost-driver for cultivated meat production, maximizing biomass per volume of media is critical to achieving efficient outcomes[32]. Additionally, despite VCD of non-adherent cells, viability percentages at equivalent times after passaging are equivalent (Fig. 1Cii). This indicates that media consumption rates, even with increased density, are not negatively impacting cell growth. These findings suggest that suspension proficiency may result in optimal culture productivity through intensifying cell densities, rather than increasing growth kinetics.

### 2. Transcriptome-Wide Response Accompanies Phenotype Transition

Due to the variation in cellular morphology and growth kinetics, we hypothesized that distinct, underlying transcriptomic profiles were responsible for driving these changes. To capture these subtle temporal changes in gene expression, RNA was collected from cells at each step in the suspension adaptation pipeline and sequenced for downstream analysis (Fig. 1A).

#### 2.1 Cells in the Adaptation Pipeline Follow a Distinct Trajectory

Principal component analysis (PCA) revealed tight clustering between biological replicates of each time point. This result indicated consistency and minimal batch effects between replicates, while clear separation was observed between time points, suggesting distinct transcriptomic changes at each point in the adaptation protocol (Fig. 2A). The first principal component (PC1) captured 56% of variance in the normalized dataset and represented variation in aggregate size. The second principal component (PC2) accounted for 29% of the variance and was interpreted as the degree of shear stress experienced by the cells. The third and fourth principal components captured 7% and 2% of the remaining variance, respectively (Supplementary Fig. 2).

**Figure 2:**
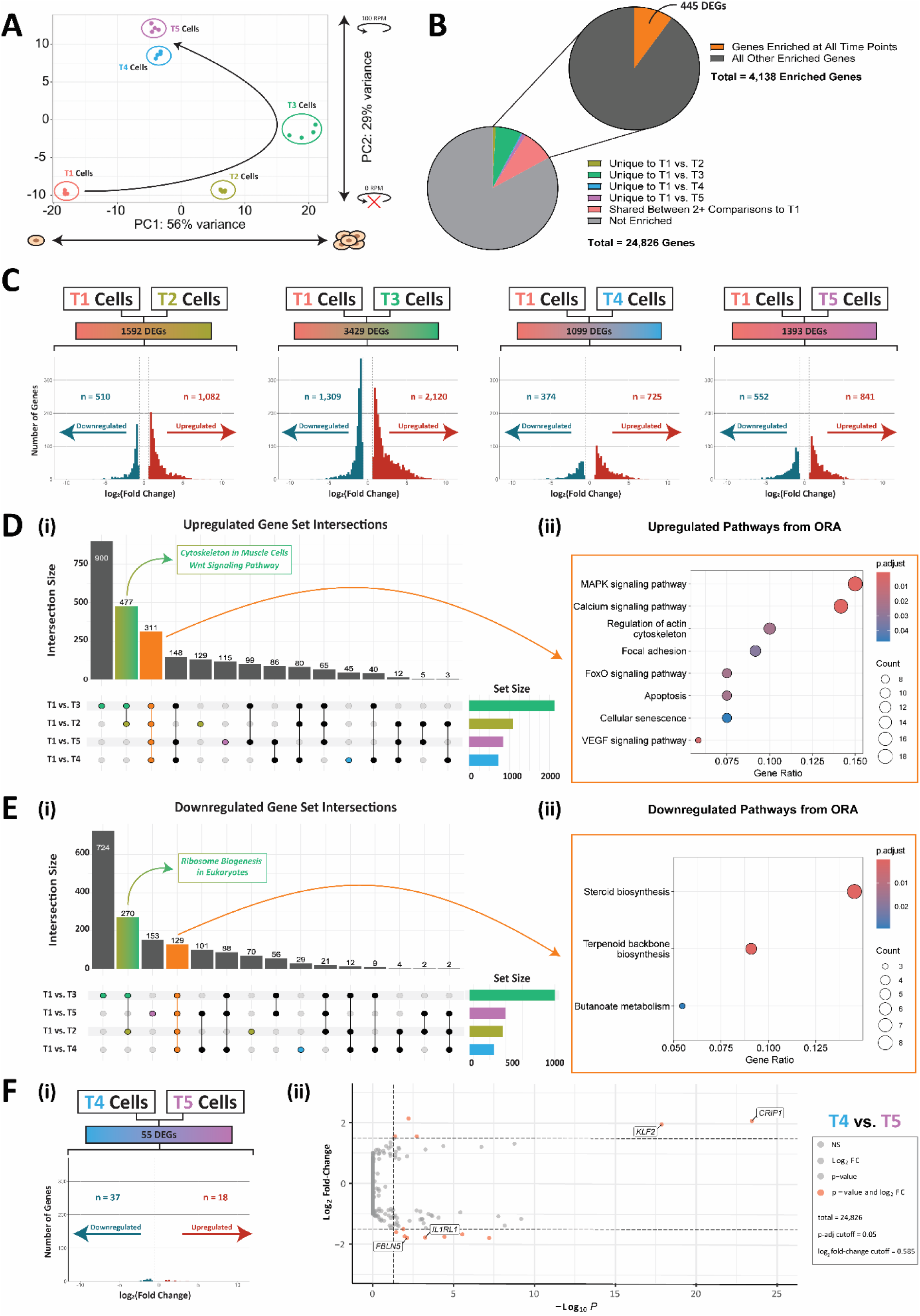
Transcriptome-wide remodeling accompanies phenotype transition. (A) PCA of DF-1 cells at each time point in the adaptation pipeline; n = 4 for each time point. PC1 represents variation in aggregate size, while PC2 represents variance in shear stress. Pink circles represent adherent T1 cells; olive circles represent T2 spheroids; green circles represent T3 aggregates; blue circles represent T4 cells at 80 RPM; and purple circles represent T5 cells at 100 RPM. Arrow represents trajectory of gene expression over time. (B) Pie chart revealing the proportion of differentially expressed genes which are unique or common to each time point contrast against T1. (C) Mirrored bar plots containing the number and log_2_ fold-change of differentially expressed genes for each time point contrast against T1. Downregulated genes are represented by blue bars, while upregulated genes are represented by red bars. Dashed lines represent the threshold value of |log_2_ fold-change| > 0.585, corresponding to 1.5-fold change. (D, i) UpSet plot representing the intersection of upregulated or (E, i) downregulated differentially expressed genes between each time point contrast against T1. The yellow-green bar represents the intersection of T1 vs. T2 and T1 vs. T3, while the orange bar represents the overlap between all contrasts. (D, ii) Pathways enriched among upregulated or (E, ii) downregulated genes from the core set of 445 genes enriched at all time points when contrasted to the adherent baseline. (F) Mirrored bar plot (i) and (ii) volcano plot representing the number of differentially expressed genes between T4 and T5 cells. Downregulated genes are represented by blue bars or circles, while upregulated genes are represented by red bars or circles. Vertical dashed lines represent the threshold value of |log_2_ fold-change| > 0.585, corresponding to 1.5-fold change. The horizontal dashed line in the volcano plot represents the adjusted p-value cutoff of 0.05.

As cells progressed through the suspension adaptation pipeline, their transcriptional profiles followed a characteristic path (Fig. 2A). Beginning as single, adherent cells during T1, aggregate size increased from T2 to T3, causing gene expression profiles to progress along PC1. As shear stress acting on T3 aggregates in static suspension culture progressively intensified during T4 and T5, transcription profiles then begin to progress along PC2. As aggregates then dissociate due to shear stress, T4 and T5 gene expression profiles regress back along PC1 while continuing along PC2. Together, the first and second principal components represent 85% of the total variance in the dataset, suggesting that both aggregate size and magnitude of shear stress drive transcriptomic changes along a predictable trajectory during the suspension adaptation process.

#### 2.2 Early Time Points in Adaptation Pipeline Represent a Critical Transition State

After conducting PCA, DGEA revealed transcriptional changes at each time point when contrasted to T1, designated as the adherent baseline (Fig. 2C). Differentially expressed genes were identified using a 1.5-fold change threshold (log_2_ fold-change > 0.585) and 5% Benjamini-Hochberg false discovery rate (BH-FDR). DGEA revealed disproportionate numbers of differentially expressed genes (DEGs) across these 4 contrasts, with early time points T2 and T3 showing the largest gene expression changes from baseline, with 1592 and 3429 DEGs, respectively (Fig. 2C). Later time points show fewer DEGs, with T4 and T5 yielding only 1099 and 1393 enriched genes, respectively, when compared to T1 (Fig. 2C). This peak in transcriptional response, comprising 1624 unique DEGs, could be attributed to the fact that at T3, cells are simultaneously downregulating adherent-required genes while upregulating genes necessary of suspension growth. This transient maximum in transcriptional changes could represent a point at which the cells commit to their new suspension phenotype.

To further investigate this hypothesis, UpSet analysis revealed that the largest intersection consisted of genes common to T2 and T3 contrasts against T1, containing 477 upregulated genes and 270 downregulated genes (Fig. 2Di, Ei). The shared genes consistently upregulated at T2 and T3 were enriched for activation of signal transduction and cell motility pathways, indicating that cells are adopting a more migratory, communicative state (Fig. 2D, ii). More specifically, the *Wnt signaling pathway* and *muscle cell-like cytoskeletal structure* are shown to be upregulated in T2 and T3 (Supplementary Table 1). Simultaneously, downregulated genes represented suppression of *ribosomal biogenesis*, possibly explaining the smaller relative size of non-adherent DF-1s (Fig. 2E, i)[33]. As these time points are midway through the adaptation pipeline, this unique overlap between T2 and T3 genes could indicate a temporary transcriptional program that subsides after adopting suspension-required genes. Coupled with the large number of DEGs obtained from the T1 against T3 contrast, this data demonstrates that the suspension adaptation process is not a simple linear progression of gradual gene expression changes.

#### 2.3 Core Set of DEGs Reveal Suppressed Lipid Metabolism, Activated Signaling and Motility

More broadly, DGEA revealed that around 17% of all genes were differentially expressed in one or more time points when compared to T1 (Fig. 2B). Of this 17%, a core set of 445 genes was enriched at all time points when contrasted to the adherent baseline (Fig. 2B, Supplementary Table 2). UpSet analysis revealed that 311 of these genes were consistently upregulated and 129 genes were consistently downregulated across all contrasts, while 5 genes did not follow a consistent trend (Fig. 2Di, Ei). Over-representation analysis (ORA) revealed that ubiquitously downregulated genes were associated with suppression of non-essential lipid metabolic pathways, such as *steroid biosynthesis* and *terpenoid backbone synthesis*, and *butanoate metabolism* (Fig. 2E, ii). In contrast, ORA of common upregulated genes showed activation of cellular processes and signal transduction pathways (Fig. 2D, ii). These include some pathways expected to change as cells transition to an anchorage-independent phenotype, such as *focal adhesion* and *regulation of actin cytoskeleton*, which both show differential expression of integrin-beta 3 (*ITGB3)*, platelet-derived growth factor receptor alpha and beta (*PDGFRA/B*), and myosin light chain 10 (*MYL10*). Notably, the transcription factor *NFAT2* was differentially expressed at all time points but implicated in the activation of multiple different pathways: *cellular senescence*, *calcium signaling*, and *VEGF signaling* (Fig. 2D, ii). Together, these central collections of genes represent a permanent shift in transcriptional state to a metabolically suppressed but dynamic, responsive, and migratory phenotype.

#### 2.4 Late-Stage Transcriptional Changes Reflect Stabilization of Non-Adherent Cells

Because PCA revealed tight clustering of T4 cells cultured at 80 RPM and T5 cells cultured at 100, a contrast between the two transcriptional states was performed to identify the stability of DF-1 cells in suspension as shear stress increased. Only 55 out of 24,826 genes (0.22%) were differentially expressed, indicating virtually no significant difference in transcriptional profiles between the two transcriptional states (Fig. 2F, i). The lack of differential gene expression over the 20-day time course (11 days at T4 and 9 days at T5) indicates transcriptional equilibrium, as gene expression has been shown to plateau within 2 hours of changing conditions[34]. Additionally, a volcano plot of this contrast revealed that DEGs in the T4 vs. T5 contrast had relatively small expression changes with log_2_ fold-changes ranging from -1.80 to 3.25 (Fig. 2F, ii). Because of the robust stability observed even with a 20 RPM increase between time points, it is likely that further time points would not yield any new, distinct transcriptional profiles. These findings indicate that slight increases in shear stress did not significantly change the expression profile of DF-1s in shaker flasks. Given that previous studies have estimated parameters of 30 to 120 RPM in spinner flasks or 80 to 100 RPM in shaker flasks for cultivated meat production, it is likely that the robustness of suspension-adapted DF-1s will be useful in bioprocessing applications[4,20,35–37].

### 3. Gene Set Enrichment Analysis Indicates Cellular Resilience After Adaptation

Beyond individual gene expression changes identified through DGEA, GSEA for each contrast was conducted using the Kyoto Encyclopedia of Genes and Genomes (KEGG) database to better understand broader functional changes[38].

#### 3.1 Acute Response is Observed Early in the Adaptation Pipeline

GSEA revealed that the T2 against T1 contrast had 45 differentially enriched pathways (DEPs), the most of any contrast against the adherent baseline (Fig. 3A). This comparison was followed by T3 with 36 DEPs, T4 with 20 DEPs, and T5 with 7 DEPs, each when contrasted against T1 (Fig. 3A). The number of DEPs decreasing over time is consistent with an acute stress response, as many of the pathways enriched at these time points imply that cells are facing challenges as they enter suspension culture. For example, the *mitogen-activated protein kinase (MAPK) signaling pathway* has been shown to mediate apoptosis, which often occurs simultaneously with *autophagy* (Fig. 3B)[39,40]. Both of these pathways are uniquely enriched in T2 and T3, marking this as an acute stress response. This is supported by suppression of *DNA replication* and repair mechanisms, such as *mismatch repair, base excision repair,* and *mRNA surveillance* in these time points exclusively, indicating commitment to cellular death or strategic resource allocation (Fig. 3B). Notably, pathways directly involved in cell junctions such as *focal adhesion* and *ECM-receptor interaction* are activated at these time points only, suggesting that the early suspension phenotype is characterized by an overexpression of cell adhesion molecules (CAMs) (Fig. 3B).

**Figure 3:**
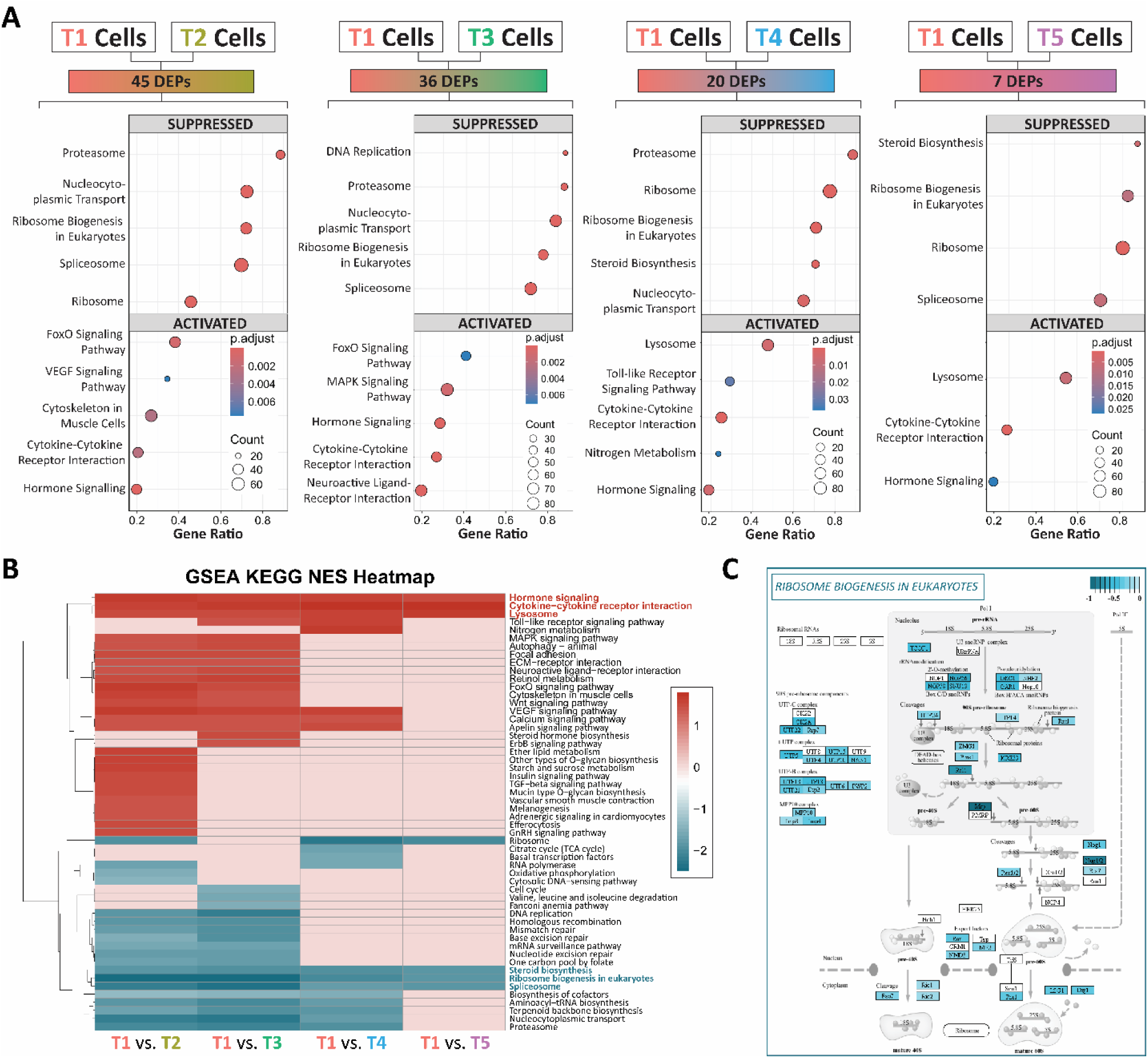
Gene set enrichment analysis of DF-1s during suspension adaptation. (A) Dot plots, ordered by gene ratio (i.e. the number DEGs implicated in a particular pathway divided by the total number of DEGs), revealing the top five most suppressed or activated pathways for each time point contrast against T1. The size of each circle represents the number of genes implicated in each pathway, while the color of the circle represents the adjusted p-value. (B) Heatmap of differentially enriched pathways observed in each time point contrast against T1. Red cells represent an activated pathway while blue cells represent a suppressed pathway. Pathway names colored red or blue indicate pathways ubiquitous across all contrasts. (C) Pathview image of the ribosome biogenesis in eukaryotes pathway, one of the commonly suppressed pathways across all contrasts against T1. Genes colored in blue are downregulated, with darker blue colors representing a higher degree of downregulation.

#### 3.2 Cells Recover from Acute Response and Return Towards Original Transcription State

Despite the stress response observed at early time points, the number of DEPs continually decreased over time despite increasing changes in cellular morphology. Importantly, many of the inflammatory and adhesion pathways implicated in this initial reaction are seen to subside by T4, indicating stabilization and cellular investment in survival. Cells at this stage re-activate repair mechanisms relating to DNA replication transcription, suggesting cellular commitment to genomic stability and mRNA quality (Fig. 3B). Though the quality of nucleic acid processes seems to recover over time, quantity diminishes, as evidenced by suppression of *basal transcription factors* and *RNA polymerase* pathways (Fig. 3B). Additionally, ATP-generating processes like *oxidative phosphorylation* and *valine, leucine, and isoleucine degradation* pathways are suppressed at T2 and T3, respectively, possibly explaining the reduction in transcription processes (Fig. 3B). Coupled with the suppression of *biosynthesis of cofactors, nucleocytoplasmic transport,* and *proteasome* pathways, cells in T4 of the pipeline are likely decreasing protein synthesis and degradation as they adapt to new conditions. However, by T5, these pathways have subsided, indicating the return of normal transcriptional and translational capabilities in the cell (Fig. 3A). Thus, the sparse number of DEPs observed in the T1 and T5 contrast demonstrates that fully adapted cells in suspension show a high degree of resiliency and recover functionality comparable to their adherent counterparts.

#### 3.3 Long-Term Cellular Phenotype is Defined by Core Set of Pathways

To better understand the permanent transcriptomic changes that occur following suspension adaptation, pathways that were enriched at all time points when contrasted to the adherent baseline were examined (Fig. 3B). A total of six pathways (three activated and three suppressed) were enriched ubiquitously across all contrasts, with these same pathways comprising 86% of all DEPs from the T1 and T5 contrast. This high degree of overlap indicates that these six pathways likely regulate the long-term phenotype differences between adherent and suspension DF-1 cells.

Of the suppressed DEPs common to all contrasts, *spliceosome* and *ribosome biogenesis* pathways represent silencing of transcription and translation machinery, respectively (Fig. 3B). Lipid metabolism was also diminished, as *steroid biosynthesis* was shown to be suppressed in all contrasts (Fig. 3B). Together, this reduction in foundational cellular machinery suggests that suspension cells exist in a lower metabolic state than adherent cells, possibly to conserve energy as cells face shear stress challenges in non-adherent conditions.

Because *ribosome biogenesis* was found to be one of the top five suppressed gene sets for all contrasts, the genes contributing to this DEP were visualized via Pathview (Fig. 3C). Overall, this analysis revealed downregulation of genes involved in all steps of ribosomal biogenesis, from early assembly and maturation to nuclear export. Expression of nearly all 90S pre-ribosomal components was reduced, which may be explained by the suppression of the RNA polymerase pathway leading to reduced levels of rRNA (Fig. 3B). Additionally, ribosomal assembly factors such as *Kre33,* maturation factors such as *Rio2,* and export factors such as *NMD3* show reduced levels of expression, indicating suppression of downstream processes as well. In total, 31.7% of all genes in this pathway were downregulated in the anchorage-independent cells, indicating an extensive, synchronized response to growth in suspension.

Activated pathways common to all contrasts were associated with environmental information processing pathways, such as signal transduction and signaling molecules (Fig. 3A). More specifically, *hormone signaling*, *cytokine-cytokine receptor interaction*, and *lysosome* pathways were activated at all time points compared to T1 (Fig. 3B). A pathway visualization of *cytokine-cytokine receptor interactions* reveals that molecules in the TGF-β family, TNF family and IL4-like class represent most of the upregulated genes in this DEP (Supplementary Fig. 3).

### 4. Soft Clustering Analysis Displays Distinct Gene Expression Patterns

To gain a broader understanding of gene expression trajectories over time, soft clustering of transcriptomic profiles was conducted via Mfuzz analysis (Fig. 4A). This algorithm is designed for transcriptomics time-series data and allows for gene assignment to multiple clusters, allowing for more robust results compared to hard clustering[41]. Additionally, utilizing time-course analysis allows for a more gene-centric approach compared to traditional GSEA.

**Figure 4:**
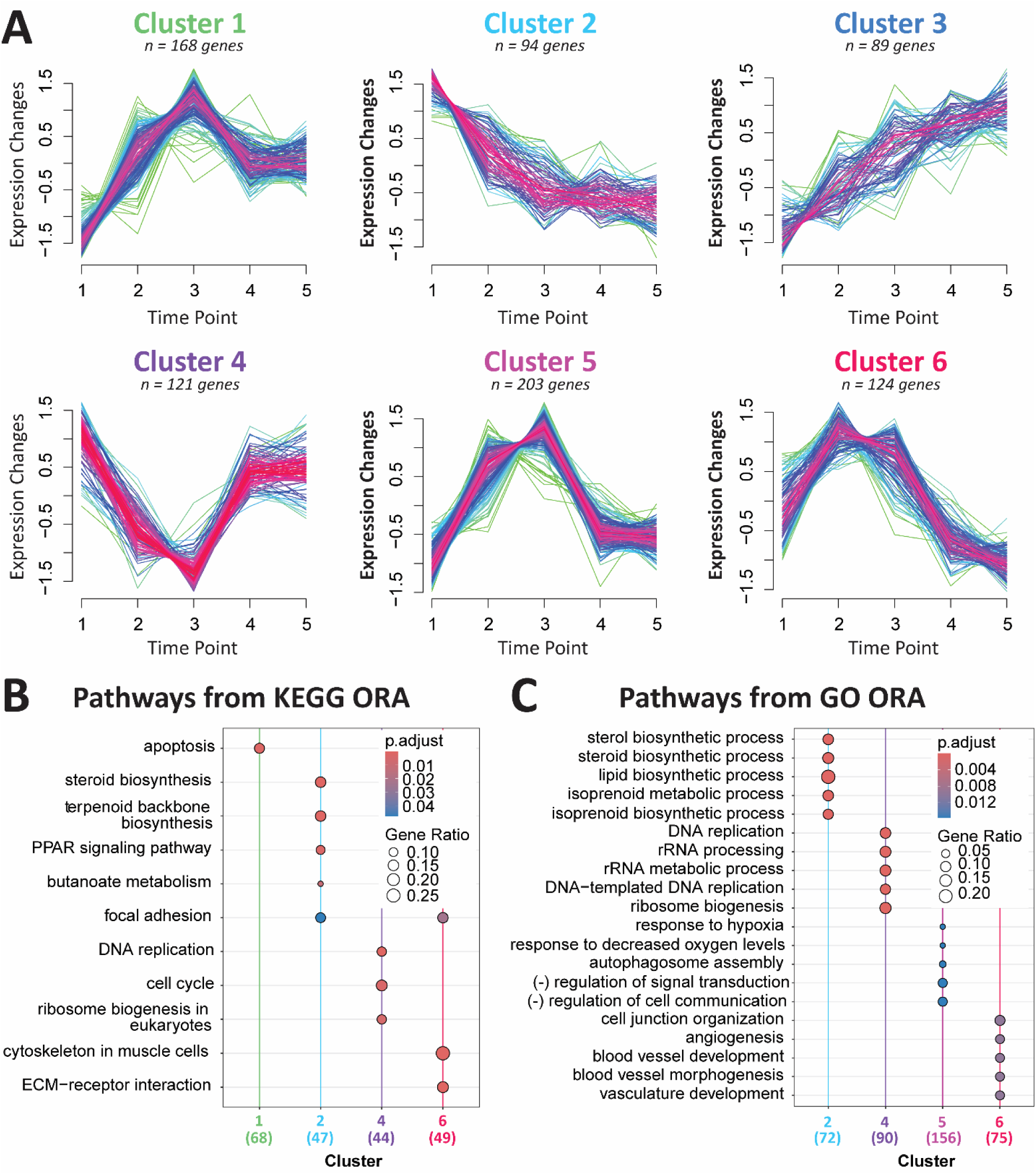
Soft clustering analysis displays distinct gene expression patterns. (A) Six clusters were generated using Mfuzz soft clustering. Individual gene trajectories are shown in multiple colors for visualization clarity only. (B) Pathways enriched for each Mfuzz cluster using the KEGG database and ORA. (C) Pathways enriched for each Mfuzz cluster using the GO database and ORA.

#### 4.1 Gene Expression Follows Six Temporal Patterns Throughout Adaptation

From this analysis, six expression patterns (i.e. clusters) were generated (Fig. 4A). Following clustering, genes within each trajectory were subjected to functional enrichment analysis to identify over-represented biological processes, enabling evaluation of pathway-level temporal dynamics. Both KEGG and Gene Ontology (GO) databases were used for enrichment of Mfuzz clusters[42,43]. Genes followed either a directional trajectory (e.g. clusters 2 and 3) or transient response pattern (e.g. clusters 1, 4 - 6), and clusters ranged in size from 89 genes to 203 genes (Fig. 4A).

#### 4.2 Transient Responses Suggest Stress and Cell Junction Remodeling

Clusters 1, 4, and 5, showed the most extreme expression changes at T3, followed by stabilization at later stages. Similar to findings from DGEA and GSEA, T3 seems to be the most stressful point in the adaptation pipeline, with gene expression peaking in Clusters 1 and 5. This peak in expression is due to cellular assembly of *autophagosomes* and preparation for *apoptosis* (Fig. 4B, C). This response seems to be due to an increase in *hypoxia*, likely due to the cells in the interior of aggregates (∼1 mm in diameter) receiving decreased levels of oxygen (Fig. 4C, Supplementary Fig. 1). Conversely, the pattern of gene expression in Cluster 4 shows a sharp drop at T3. This decrease in expression of *DNA replication* and *ribosome biogenesis* could be caused by the hypoxic stress cells are simultaneously experiencing, forcing the DF-1s to allocate resources towards survival rather than gene expression (Fig. 4B, C).

The trajectory observed from Cluster 6 instead showed a gene expression maximum occurring at T2 with later downregulation of genes. At this point, cells are overexpressing genes involved with changes to *cell junctions,* which comprise the connection between two cells or between a cell and the extracellular matrix (Fig. 4B, C)[44]. More specifically, *focal adhesion* and *ECM-receptor interaction* pathways are activated based on genes in the Cluster 6 trajectory. Because T2 spheroid size is controlled by the geometry of the plate, cells with sufficient oxygen can instead direct energy reserves towards adjusting to an environment without attachment surfaces. Additionally, enrichment with the GO database showed that *angiogenesis* and *vasculature* pathways are activated at this time point, potentially implying that cells are signaling for blood vessel formation to prevent eventual hypoxia.

#### 4.3 Sustained Responses Show Permanent Changes in ECM Remodeling and Apoptosis

Cluster 2, consisting of 94 genes, displayed synchronous downregulation of genes and was enriched for *lipid* and *steroid biosynthesis* pathways from both GO and KEGG databases (Fig. 4B, C). As discovered by DGEA and GSEA earlier in this work, cluster 2 corroborates the idea that suspension cells have reduced lipid metabolism compared to adherent cells. Genes belonging to the *focal adhesion* pathway also followed the downward trajectory in cluster 2, indicating that CAMs may decrease over time (Fig. 4B). Additionally, *PPAR signaling* pathway genes fall in this cluster, which may have important implications for the transdifferentiation potential of DF-1 cells into adipocytes (Fig. 4B)[4,45,46].

Conversely, Cluster 3 showed consistent upregulation of 89 genes, but had no significant enrichment in KEGG or GO databases (Fig. 4B, C). Although these uncharacterized genes lacked common annotations, some common functional themes emerged with manual examination (Supplementary Table 3). For example, ECM-remodeling and/or pro-angiogenic genes fall into this cluster, such as *EFEMP1, ECM1,* and *COL26A1*[47–49]. Similarly, genes like *PDLIM1, FAM129A,* and *SDK2* show cytoskeletal and CAM re-organization[50–52]. Notably, Cluster 3 genes *PHLDA1* and *OSGIN1* have both been shown to induce p53-dependent apoptosis, potentially indicating that suspension cells are primed to eliminate unstable cells which cannot survive the adaptation process[53,54].

### 5. Validation of Differentially Expressed Genes by qPCR

In order to confirm the RNA-seq analysis results, quantitative real-time polymerase chain reaction (qPCR) was conducted on differentially expressed genes identified through transcriptomic analysis. A subset of 11 genes was chosen primarily based on their adjusted p-value in the T1 vs. T5 contrast, and an expression profile of log_2_ fold-change of greater than 1 or less than -1. Three of these genes, cadherin EGF LAG seven-pass G-type receptor (*CELSR1),* claudin 11 (*CLDN11),* and collagen type XII alpha 1 chain *(COL12A1),* were selected based on biological relevance to cell adhesion[55–57].

#### 5.1 Gene Expression Results from qPCR Correlate with Results from RNA-Seq

Directionality of gene expression was conserved between both methods, confirming changes in transcript regulation (Fig. 5A). Additionally, the relative expression of the selected genes from qPCR and RNA-seq was significantly correlated with a Pearson correlation coefficient of 0.8586 (p < 0.001) (Fig. 5B). Overall, the uniformity in the magnitude and direction of differential gene expression across methods suggests reliable results from the RNA-seq analysis.

**Figure 5:**
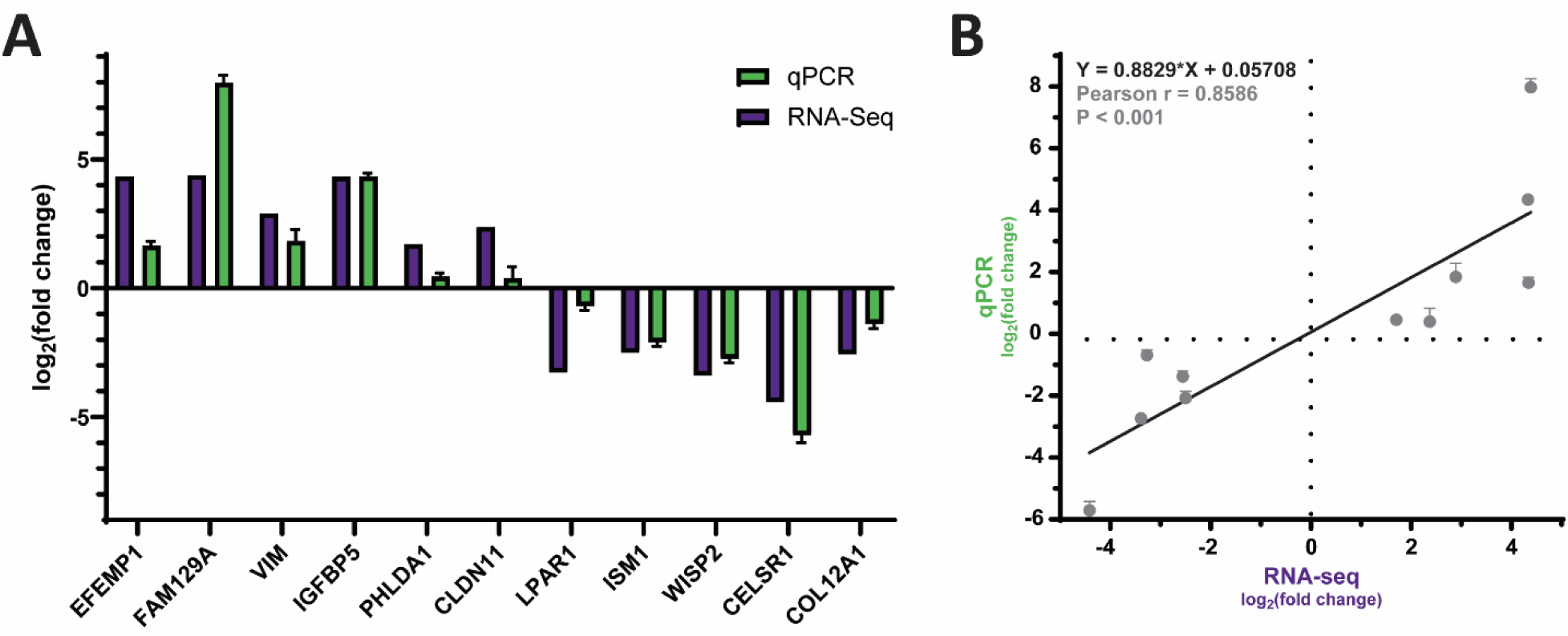
Validation of differentially expressed genes by qPCR. (A) The log_2_ fold-change results from qPCR for differentially expressed genes compared to those generated from RNA-seq analysis. Of the differentially expressed genes, 3 were chosen for biological relevance (*CLDN11, CELSR1, COL12A1)* and 8 were chosen for statistical significance. Error bars represent standard deviation; n = 6. (B) Linear regression analysis of the log_2_ fold-change values between RNA-seq analysis and qPCR. Error bars represent standard deviation; Pearson correlation coefficient (r) = 0.8586; p-value < 0.001; n = 6.

## Discussion

For cultivated meat to continue to progress as a technology, “domesticating” cells to possess phenotypes suitable for scaled-up, cost-efficient bioproduction is required. This need includes reduced adherence and anoikis resistance[58]. Suspension cells with these characteristics are able to achieve significantly higher VCDs compared to both traditional adherent culture and microcarriers, enabling the production of more cultivated biomass[59]. However, the inherent anchorage-dependent nature of many food-relevant cell lines has limited the widespread adoption of suspension culture for cultivated meat bioprocesses. The understanding of the mechanisms that occur during the transition of food-relevant cell lines to suspension provides the foundation for the development of future cell lines capable of non-adherent growth.

In the present work, we successfully adapted adherent DF-1 cells to suspension culture and performed transcriptomic analysis on the cells at each stage in the granular adaptation pipeline. Using DGEA, GSEA, and Mfuzz clustering, we elucidated a timeline of gene expression as the cells adopted a suspension-proficient phenotype (Table 1). Notably, we found that gene and pathway dynamics could be grouped based on their temporal gene expression into either transient, sustained, or progressive responses, with virtually no late-onset responses. This could indicate that DF-1s exhibit a hormetic response upon entering suspension, causing cells to implement adaptive compensatory pathways in order to survive new culture conditions[60].

**Table 1:** Summary of temporal dynamics as DF-1 cells transition from adherent to suspension.

| Theme | Pathway | Expression Relative to T1 |  |  |  | Method / Evidence | Dynamics | Biological Significance |
| --- | --- | --- | --- | --- | --- | --- | --- | --- |
|  |  | Early | Mid | Late | Pattern |  |  |  |
| Response to Stress | Autophagy | ↑↑ | ↑↑↑ | → | peak at T3, return to baseline | GSEA (T2, T3)<br>Mfuzz (Cluster 5) | Transient Activation | Cells experience increased autophagy at T2 and T3 that subsides, but apoptosis and senescence are sustained in suspension cell populations compared to adherent populations. |
|  | Apoptosis | ↑↑ | ↑↑↑ | ↑ | peak at T3, sustained upregulation | DGEA (445 core genes)<br>Mfuzz (Cluster 1) | Sustained Activation |  |
|  | Cell Senescence | ↑ | ↑ | ↑ | sustained activation | DGEA (445 core genes) | Sustained Activation |  |
| Adhesion | Focal Adhesion | ↑↑↑ | ↑↑ | ↓↓↓ | peak at T2, trough at T5 | DGEA (445 core genes)<br>GSEA (T2, T3)<br>Mfuzz (Cluster 2, 6) | Transient Activation, Sustained Suppression | As a response to losing attachment surfaces, cells initially upregulate CAMs in a desperate attempt to maintain adhesion. However, after adapting to suspension culture, cells have lower levels of CAMs than adherent counterparts. |
|  | ECM-Receptor Interaction | ↑↑↑ | ↑↑ | ↓↓↓ | peak at T2, return to baseline | GSEA (T2, T3)<br>Mfuzz (Cluster 6) | Transient Activation, Sustained Suppression |  |
|  | Cell Junction Organization | ↑↑ | ↑↑↑ | → | peak at T3, return to baseline | Mfuzz (Cluster 5) | Transient Activation |  |
| Cytoskeleton & Structure | Regulation of Actin Cytoskeleton | ↑ | ↑ | ↑ | sustained activation | DGEA (445 core genes) | Sustained Activation | As actin is directly related to cell adhesion, regulation of cytoskeletal proteins is upregulated at all time points with a temporary upregulation of muscle cytoskeletal proteins at T2 and T3. |
|  | Cytoskeleton in Muscle Cells | ↑↑↑ | ↑↑ | ↓↓↓ | peak at T2, trough at T5 | DGEA (T2, T3 intersection)<br>GSEA (T2, T3)<br>Mfuzz (Cluster 6) | Transient Activation |  |
| Nucleic Acid Processing | Ribosome Biogenesis in Eukaryotes | ↓↓ | ↓↓↓ | → | trough at T3, sustained downregulation | DGEA (T2, T3 intersection)<br>GSEA (T2, T3, T4, T5)<br>Mfuzz (Cluster 4) | Sustained Suppression | Sustained downregulated nucleic acid processing at virtually all time points indicates that cells are less metabolically active than adherent cells, due to both size and stress from being in suspension culture. |
|  | DNA Replication | ↓↓ | ↓↓↓ | → | trough at T3, return to baseline | GSEA (T2, T3)<br>Mfuzz (Cluster 4) | Transient Activation |  |
|  | Spliceosome | ↓ | ↓ | ↓ | sustained downregulation | GSEA (T2, T3, T4, T5) | Sustained Suppression |  |
| Bioactive Lipid Metabolism | Steroid Biosynthesis | ↓ | ↓↓ | ↓↓↓ | sustained downregulation | DGEA (445 core genes)<br>GSEA (T2, T3, T4, T5)<br>Mfuzz (Cluster 2) | Progressive Suppression | Progressively suppressed cell metabolism pathways in suspension cells may be due to stress, decreased cell size over time, hypoxia, changes in morphology or some combination of these factors. |
|  | Terpenoid Backbone Biosynthesis | ↓ | ↓↓ | ↓↓↓ | sustained downregulation | DGEA (445 core genes)<br>GSEA (T2, T3, T4)<br>Mfuzz (Cluster 2) | Progressive Suppression |  |
|  | Butanoate Metabolism | ↓ | ↓↓ | ↓↓↓ | sustained downregulation | DGEA (445 core genes)<br>Mfuzz (Cluster 2) | Progressive Suppression |  |
| Cellular Communication | MAPK Signaling | ↑ | ↑↑ | → | peak at T2 or T3, sustained upregulation | DGEA (445 core genes)<br>GSEA (T2, T3) | Sustained Activation | Suspension culture causes DF-1s to adopt a more communicative state compared to adherent counterparts, though negative regulation of signal transduction attempts to lessen this activation at early stages. Specifically, these signaling pathways represent stress responses and efforts to return to adherent expression patterns. |
|  | Calcium Signaling | ↑ | ↑↑ | → | peak at T2 or T3, sustained upregulation | DGEA (445 core genes)<br>GSEA (T2, T3, T4) | Sustained Activation |  |
|  | FoxO Signaling | ↑ | ↑↑ | → | peak at T2 or T3, sustained upregulation | DGEA (445 core genes)<br>GSEA (T2, T3) | Sustained Activation |  |
|  | Hormone Signaling | ↑ | ↑ | ↑ | sustained upregulation | GSEA (T2, T3, T4, T5) | Sustained Activation |  |
|  | Cytokine-Cytokine Receptor Interaction | ↑ | ↑ | ↑ | sustained upregulation | GSEA (T2, T3, T4, T5) | Sustained Activation |  |
|  | Negative Regulation of Signal Transduction | ↑↑ | ↑↑↑ | → | peak at T3, return to baseline | Mfuzz (Cluster 5) | Transient Activation |  |
| Angiogenesis | VEGF Signaling | ↑ | ↑ | → | sustained upregulation | DGEA (445 core genes)<br>GSEA (T2, T3, T4) | Sustained Activation | Fibroblasts in suspension are more senescent than adherent counterparts, causing VEGF expression. However, hypoxia plays a role in VEGF expression at early stages. |
|  | Vasculature Development | ↑↑↑ | ↑↑ | ↓↓↓ | peak at T2, trough at T5 | Mfuzz (Cluster 6) | Transient Activation |  |

As cells transition through the adaptation pipeline, we hypothesize that two environmental stressors, oxidative stress and cell detachment, drive transcriptomic changes (Fig. 6A)[18]. Hypoxia, a form of oxidative stress, is known to impact spheroids larger than around 200 μm due to limited oxygen diffusion from the environment to the core[61]. Because cellular aggregation occurs only at T2 and T3, hypoxia-driven responses are transient and resolve after these time points. For example, GSEA and Mfuzz analysis indicate that cells at these points attempt to acutely conserve resources through *autophagy* and halting *DNA replication* (Fig. 3B, 4B, 6A)[62]. Hypoxia has previously been shown to drive decreases in DNA repair, which was observed in GSEA, possibly explaining the inhibition of DNA synthesis (Fig. 3B)[63]. Similarly, Mfuzz captured that *signal transduction* and *cellular communication* are negatively regulated early in the pipeline, which is likely caused by efforts to prevent excess resource usage, desensitization, or overreaction (Fig. 4C)[64]. In addition, all three analysis methods revealed transient *muscle cell-like cytoskeletal remodeling*, alongside activation of *VEGF signaling* and *vasculature development* pathways, suggesting a pro-angiogenic response to hypoxic conditions (Table 1)[65]. Though the VEGF signaling pathway remained activated at later stages, it is likely that this response was due to a combination of sustained *cellular senescence* rather than hypoxia at later stages (Fig. 2Dii)[66]. Together, the transient dysregulation of these pathways represent mechanisms that are not necessary for long-term survival in suspension but may be an important step in the adaptation process.

**Figure 6:**
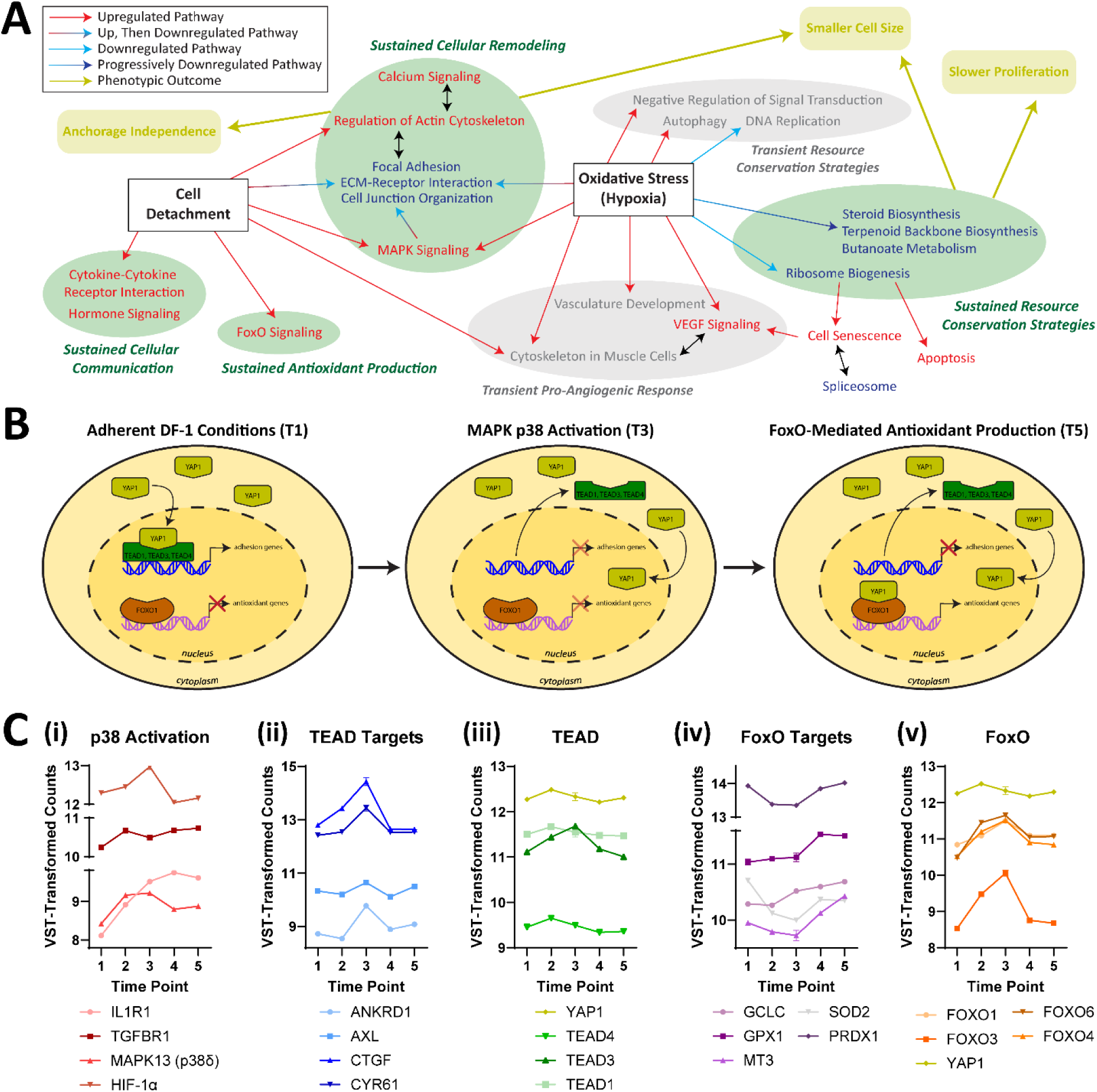
Proposed mechanism of suspension proficiency in DF-1 cells. (A) Bubble map showing the relationship between enriched pathways as DF-1s transition from adherent to suspension conditions. Different groupings of enriched pathways represent various cellular responses, which in turn lead to phenotypic outcomes. (B) Schematic of proposed mechanism DF-1s utilize to survive the suspension adaptation. (C) VST-transformed RNA-seq counts of (i) p38-relevant genes, (ii) TEAD target genes, (iii) TEAD itself, (iv) FoxO target genes, and (v) FoxO itself. Error bars represent standard error, n = 4.

Despite hypoxia only existing in early time points, the effects of oxidative stress seem to influence transcription patterns beyond early stages, evidenced by a peak in expression which plateaus at a new steady state for the remainder of the adaptation process. For example, cells attempt to conserve resources through the progressive suppression of non-essential lipid metabolism, such as *steroid biosynthesis* (Table 1, Fig. 6A). Importantly, cells also downregulate *ribosomal biogenesis,* one of the cell’s most resource-intensive processes, in response to hypoxia-driven nucleolar disruption (Table 1)[67]. The suppression of ribosomal synthesis results in increased *cellular senescence* and *apoptosis,* but also may explain the reduction in cell size and growth rate of suspension cell lines (Fig. 6A)[67,68]. Previous work has demonstrated that downregulation of ribosomal biogenesis leads to smaller cell growth (i.e. size), which in turn inhibits cell proliferation (i.e. doubling); both changes were observed in suspension cells compared to adherent counterparts (Fig. 1Ci, iii)[69,70]. Though these transcriptional programs initially happen concurrently with autophagy and other acute hypoxic responses, non-essential lipid metabolism and ribosome biogenesis remain suppressed and describe the cell’s long-term resource conservation strategies.

Together, oxidative stress and cell detachment work to drive some of the most expected and sustained transcriptional changes, such as shifts in CAM expression, as cells adapt to suspension. For example, hypoxia has been shown to increase the number of focal contacts and disrupt the actin cytoskeleton in fibroblasts[71]. This temporary upregulation of CAMs, a desperate attempt to restore cell-matrix contact, can be observed in Mfuzz clustering (Fig. 4B). By T4, cells have suppressed focal adhesion pathways to below adherent baseline expression, marking sustained downregulation of CAMs (Fig. 4B). Because the cytoskeleton and CAMs are directly linked, a consistent upregulation in *calcium signaling* likely allows the cell to *regulate the actin cytoskeleton,* thus controlling *cell junction organization* (Table 1)[15,72–75]. This behavior suggests a hormetic response, with cells shifting their energy towards novel survival strategies rather than reverting to the adherent phenotype.

In addition, GSEA identified general activated pathways that indicate increased cellular crosstalk via cytokines and hormones, consistent with previous studies demonstrating that cells in suspension display enrichment of “immune related genes” compared to their adherent counterparts (Fig. 6A)[18]. This upregulation of communication signals reveals that suspension cells are utilizing signaling cues to permanently alter cell junctions, which has been shown in prior literature[76,77].

To this point, we propose a novel mechanism for suspension based on work from several others (Fig. 6B)[18,78,79]. Because DF-1s upregulate MAPK in response to non-adherent culture, we investigated the transcriptional signatures to determine the predominant pathway and found activation of both the classical ERK pathway and the oxidative stress-induced p38 pathway (Supplementary Fig. 4)[80]. With *HIF-1*α driving hypoxic responses*, IL1R1* and *TGFBR1* were upregulated at all time points after T1, causing a peak in p38δ at T3 before plateauing above baseline (Fig. 6Ci). We hypothesize that this p38δ activation causes the TEA domain family of transcription factors (TEADs) to translocate to the nucleus, preventing DNA-binding and interaction with yes-associated protein 1 (YAP) based on prior work (Fig. 6B)[78]. In 2023, engineering HEK293A cells with certain transcription factors disabled functionality of the YAP-TEAD transcriptional complex, which caused downregulation of adhesion genes and thus, spontaneous cell-matrix dissociation[18]. Particularly, the authors found that expression of TEAD2 was significantly decreased in engineered suspension cells[18]. Because we speculated that the translocation of TEAD prevented expression of YAP-TEAD induced adhesion genes in DF-1 cells, we examined classical TEAD target genes: *ANKRD1, AXL, CTGF,* and *CYR61* (Fig. 6Cii)[81]. We found that TEAD target genes spiked at T3 before dropping back to baseline levels during T4 and T5 despite relatively constant expression levels of TEAD, indicating that cytoplasmic transportation may lead to adhesion gene silencing as p38δ is activated (Fig. 6Cii, iii). With its preferential binding partner TEAD no longer in the nucleus, we proposed that YAP may instead interact with the transcription factor forkhead box, class O protein 1 (FOXO1), based on prior work[79]. The authors showed that FOXO1 forms a functional complex with YAP in cardiomyocytes under oxidative stress, causing downstream activation of catalase and *SOD2*[79]. Though DF-1s do not express catalase at any point in the adaptation process, we observed that expression of *SOD2* activates once TEAD had translocated at T3, supporting this hypothesis (Fig. 6Civ). Similarly, antioxidant defense genes known to be FOXO targets were upregulated only at later time points (Fig. 6Civ). Notably, FOXO expression peaked at T3 and stabilized consistently elevated above baseline levels, possibly indicating a stress response (Fig. 6Cv). Thus, we propose that in DF-1 cells, MAPK p38 activation causes TEAD sequestration that redirects YAP to FOXO1, causing simultaneous downregulation of adhesion genes and production of antioxidants. As the presence and functionality of *TEAD2* is unclear in *Gallus gallus*, DF-1s may be predisposed to activating this mechanism to gain suspension-proficiency[82].

Though these findings have similarities with other studies which investigated the difference between adherent and non-adherent cell lines, the translatability of this work is uncertain. For example, the overall downregulation of CAM-related genes was observed in anchorage-independent CHO[28] and human MSCs[83], while the opposite was seen in suspension Vero cells[84]. Paradoxically, contradictory expression patterns were observed for adhesion-related genes across different studies utilizing HEK293A[14,18]. A literature review of comparative studies between adherent and non-adherent cells found that no individual genes or gene sets function as the universal driver of suspension proficiency. However, common themes did emerge for anchorage-independent phenotypes regardless of mesenchymal or epithelial cell origin. For example, altered cytoskeletal organization and cell-matrix adhesion were observed in BHK, HEK293, Vero, MDCK, and CHO cells[14,15,17,84,85]. Similarly, changes to lipid metabolism were observed in HEK293, Vero, and human epithelial breast adenocarcinoma (MDA-MB-468)[14,84,86]. DF-1 cells and HEK293 also displayed reduced oxidative phosphorylation and nucleotide metabolism but increased antioxidant activity, possibly due to their similar embryonic origin[14,18]. Though these themes likely contribute to suspension adaptation, the specific determinants involving cell type, origin, species, and adaptation method remain unresolved for many food-relevant cell lines.

Future work should address the limitations of this study, such as repeating this work in animal-component free (ACF) culture media, a necessity for the cultivated meat bioprocess. As serum is known to protect cells and provide excess nutrients in challenging environments like suspension culture, the findings of this work may not apply to cells cultured in ACF medium[87].

Additionally, broadening the generalizability of these findings could be achieved by extending this study to multiple food-relevant species and cell types. For example, the present findings may be specific to embryonic fibroblast cells from any species, or any cell type derived from *Gallus gallus*. Most importantly, subsequent research should validate the proposed mechanism of suspension adaptation. This may be achieved by immunofluorescence to visualize TEAD in the cytoplasm after MAPK activation, co-immunoprecipitation of FOXO1 and YAP proteins, or cell-based luciferase assays to quantitatively assess YAP-FOXO1 and YAP-TEAD targets. Finally, once validated, functional studies implementing this mechanism across other cell types could enable the engineering of suspension cell lines for cultivated meat production.

## Interests

The authors declare no competing interests.

## Data and Code Availability

The authors declare that data supporting this study are available within Supplementary Information. Extra data and analysis are available from the corresponding author upon request. Raw RNA-seq files are available at the NCBI Sequence Read Archive at this link.

## Author Contributions

E.J.C contributed to methodology, conducting experiments, performing data analysis, creating figures, and writing the manuscript. A.N. contributed to methodology and performing data analysis. B.H.B. contributed to methodology and editing the manuscript. M.P.V. contributed to conducting experiments and editing the manuscript. D.L.K contributed to experimental design and manuscript drafting. All the authors have read and contributed to manuscript editing.

## Acknowledgements

We thank the United States Department of Agriculture (2021-05678) for their support of this work. The authors acknowledge the Tufts University High Performance Compute Cluster (https://it.tufts.edu/high-performance-computing) which was utilized for the research reported in this paper. We thank Kirsten Trinidad for initial ideation and guidance. We are grateful to Emily Lew, Leslie Boc, Auggie Wirasaputra, and Tianna Edwards for assistance with cell culture and qPCR. We also thank Camilo Riquelme-Guzmán and Juan Aguilera Moreno for guidance with molecular biology techniques and discussions. Finally, thank you to the Tufts University Center for Cellular Agriculture past and present members for scientific input and support. Figures were created using Adobe Illustrate 2021.

## Methods

### Cell Culture

UMNSAH/DF-1 (ATCC; CRL-12203) adherent cells were obtained from the American Type Culture Collection (ATCC, Manassas, Virginia, USA). Cells were cultured in serum-supplemented media (SSM) consisting of Dulbecco’s modified eagle medium + GlutaMAX (Gibco, Waltham, Massachusetts, USA;10569-010) supplemented with 10% fetal bovine serum (Gibco, A56707-01) and 1% antibiotic-antimycotic (Gibco; 15240-062) at 5% carbon dioxide and 39°C in a humidified incubator (ThermoFisher, Waltham, Massachusetts, USA; 51030994). Adherent DF-1s were thawed from cryopreservation at passage X+12. During routine passaging of adherent cells, cells were seeded between 2,500 – 6,000 cells/cm^2^ in 75 cm^2^ T-flasks (ThermoFisher; 156499) and grown until 80 - 100% confluency before being enzymatically dissociated using 0.25% trypsin-EDTA (Gibco; 25200-072). Cell counts, viability, and diameter were tracked using an automated cell counter (Chemometec, Copenhagen, Denmark; NucleoCounter® NC-200™). The doubling time of adherent and non-adherent cells was calculated using the following equations:

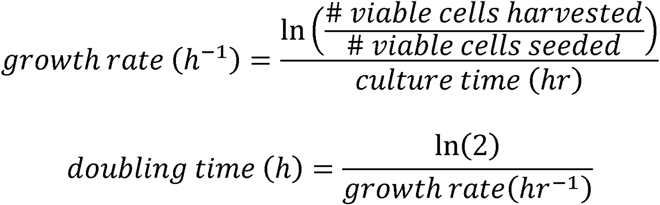

### Imaging

Cells were imaged using phase contrast microscopy on an inverted microscope (Olympus Corporation, Tokyo, Japan; CKX53). Images were capture through EPview™, an application used to stream high-resolution images from the microscope to mobile devices.

### Adaptation to Suspension

After ∼2 months (passage X+27) in adherent culture, the cells were gradually adapted to suspension culture using published methods[4]. Briefly, adherent cells were dissociated from tissue culture plastic using TrypLE (ThermoFisher; 12604021). Around 10% of these “T1” cells were lysed for RNA extraction, while the remaining 90% were plated at 1.2 million cells per well in an AggreWell™ 400 plate (STEMCELL Technologies, Vancouver, BC, Canada; 34411) pre-treated with Anti-Adherence Rinsing Solution (STEMCELL Technologies; 07010). After 24 hours of incubation, spheroids were collected using a large 37 μm strainer (STEMCELL Technologies; 27250) according to manufacturer’s instructions. Around 5% of these “T2” spheroids were lysed for RNA extraction, while the remaining 95% were seeded in ultra-low attachment T75 flask (Corning, Corning, New York, USA; 3814) and mechanically disturbed via vigorous pipetting once per day for the next 4 days. From this point forward, SSM was supplemented with 0.1% Poloxamer 188 (Gibco; 24040032) to support suspension growth. Around 5% of the “T3” cell suspension was lysed for RNA extraction, while the remaining 95% was seeded into 125 mL baffled shaker flasks (Fisher Scientific, Waltham, Massachusetts, USA; BBV12-5) and cultured in a humidified shaking incubator (Millipore Sigma, Burlington, MA, USA; Z742446-1EA) at 80 RPM, 39 °C, and 5% carbon dioxide. Twice per week, cells were collected, dissociated with TrypLE, and reseeded at 200k cells/mL. After 9 days at 80 RPM, around 5% of the “T4” cells were lysed for RNA extraction while the remaining cells were reseeded as previously described. The shaking speed was increased to 100 RPM and passaged twice per week in the same way as the T4 cells. Cells were cultured at 100 RPM for 11 days, after which 5% of the “T5” cells were lysed for RNA extraction.

### Bulk RNA Sequencing (RNA-seq) Analysis

#### Library Preparation and Sequencing

Four independent batches (n = 4) of anchorage-dependent cells were adapted to suspension for RNA sequencing analysis. Cell lysate from each time point was stored at –80 °C until all samples had been collected. RNA was extracted from simultaneously from all samples to minimize batch effects. RNA was isolated from cell lysate and purified from DF-1s at the previously specified time points using RNeasy Mini Kit (Qiagen, Hilden, Germany; 74104) according to manufacturer’s instructions. Adherent cells were dissociated from tissue culture plastic using TrypLE to avoid RNA degradation before purification[88]. RNA concentration, 260/280 ratios, and 260/230 ratios were determined using NanoDrop™ 2000 Spectrophotometer (ThermoFisher, ND-2000).

RNA samples were sent on dry ice to Novogene (Sacramento, California, USA) for library preparation and RNA sequencing. Sample quality of RNA was determined by Novogene using gel electrophoresis, Nanodrop, and Agilent2100 to determine RNA Integrating Number (RIN). Non-directional library preparation was performed and libraries were sequenced on an Illumina NovaSeq X Plus using a 25B flowcell with paired-end, 150 base pair (bp) reads.

#### Read Alignment and Quantification

Alignment, quantification, and analysis was performed using a development version of the tucca-rna-seq workflow (Git commit 571a7687a4fde40c4705cd12dbaf1250688565ff). The first stable, citable release of the software (v0.9.0) is formally cited in the references. Unless noted otherwise, for all software used, all parameters not mentioned specifically were set to the default values of the tucca-rna-seq workflow. The NCBI RefSeq assembly of the Gallus gallus reference genome (bGalGal1.mat.broiler.GRCg7b; GCF_016699485.2) and its corresponding annotation were retrieved using ncbi-datasets-cli (v16.17.1)[89]. This genome served as the reference for STAR (v2.7.11b) alignment and as the source for decoy sequences when building the Salmon (v1.10.3) decoy-aware transcriptome index[90,91].

Via the tucca-rna-seq workflow, quality control (QC) was performed for raw reads using FastQC (v0.12.1)[92]. Following alignment with STAR, Qualimap (v2.3) was used to analyze read alignment QC, determine the genomic origin of reads, and assess transcript coverage[93,94]. Transcript abundance was quantified using Salmon’s selective alignment mapping strategy, which mitigates spurious read mapping through the use of a decoy-aware transcriptome index[95]. Informed by biases identified during alignment quality control, this pseudo-alignment process also corrected for sequence-specific and positional biases by enabling the --seqBias and --posBias flags in Salmon. An HTML report summarizing the raw read QC, read alignment QC, and transcript quantification results was generated using MultiQC (v1.25) for multi-sample comparison[96].

#### Differential Expression Analysis

Transcript quantification files from Salmon were imported into R (v4.4.1; R Core Team 2024) and summarized to gene-level counts using the tximport package (v1.34.0)[97,98]. To account for potential changes in transcript length across samples, gene-level counts were generated using the lengthScaledTPM method, which creates integer-scale counts by scaling transcript-per-million (TPM) values by the library size.

The resulting list object from tximport was used to create a DESeqDataSet object. For most visualization and clustering applications, the DESeqDataSet object was transformed using the Regularized Log Transformation (rlog) function within the DESeq2 package (v1.46.0)[99]. Sample-level QC was subsequently performed on the log2 transformed normalized counts. Using DESeq2, principal component analysis was performed using singular value decomposition on the 1000 most variable genes to examine the co-variances between samples. To further explore the co-variances between samples, unsupervised hierarchical clustering was performed using the pheatmap CRAN package (v1.0.13) with complete-linkage analysis and a Euclidean distance measure[100]. No samples were identified as outliers, thus all samples were included in analysis.

Differential expression analysis was performed with DESeq2 by applying the DESeq function to the DESeqDataSet object. This method internally corrects for library size differences using the median-of-ratios method, estimates gene-wise dispersions, and fits a negative binomial generalized linear model. The DESeqDataSet object was created using a design of ∼ timepoint, where “timepoint” was a factor with two levels: T1 vs. TX. Wald tests were performed to test for temporal differences by assessing the expression of genes in T1 (adherent baseline) relative to TX (later time points). Log_2_ fold-change results were shrunken toward zero for genes with low counts and/or high dispersion values using the ashr method[101]. Genes were considered significantly differentially expressed if they exhibited a Benjamini-Hochberg False-Discovery Rate (BH-FDR) < 0.05 and an absolute log_2_ fold change > 0.585, corresponding to a > 1.5-fold change.

Principal component analysis was conducted using regularized log transformed psuedocounts, to prevent any bias from the abundance of low-count genes. Pairwise contrasts were set with T1 as the base level in all comparisons besides T4 vs. T5, where T4 was designated as the base level. The log_2_ fold-change threshold was set to 0.585 and BH-FDR (i.e. adjusted p-value) of 0.05. The organism was specified as *Gallus gallus* when applicable.

Data frames containing DEGs, log_2_ fold-change, and adjusted p-value for each contrast were obtained from DGEA and used for visualizations. For example, genes were binned by log_2_ fold-change direction and magnitude, then used to make mirrored bar plots using the ggplot2 R package (v3.5.2)[102]. UpSet plots were generated by using concatenated lists of either upregulated genes or downregulated genes from each contrast and the ComplexUpSet R package (v1.3.3)[103]. For the T4 and T5 contrast, volcano plots were produced using the EnhancedVolcano R package (v1.13.2) and previously specified FDR and fold-change cutoff[104].

#### Pathway Expression Analysis

To understand the functional implications of these expression changes, two complementary pathway analyses were performed using the clusterProfiler Bioconductor package (v4.14.6)[105,106]. Gene sets for *Gallus gallus* were used for all functional analyses; KEGG pathways were accessed using the ‘gga’ organism identifier. First, an over-representation analysis (ORA) was conducted using the list of significantly differentially expressed genes (DEGs) to identify enrichment in KEGG pathways. A background set of all genes detected in the expression matrix was used for the ORA. Second, to detect subtle, coordinated pathway-level shifts, Gene Set Enrichment Analysis (GSEA) was performed. For GSEA, all genes were pre-ranked by their shrunken log_2_ fold change values, and this ranked list was tested for enrichment against the KEGG gene set, using a seed of “12345” for reproducibility. For all pathway analyses, we identified significantly enriched terms using a BH-FDR significance level of 0.1.

The top 5 most enriched gene sets for each contrast were visualized using the dotplot() function from the clusterProfiler R package (v4.14.6). Dot plots were customized by setting x = “geneRatio” to show gene ratios (i.e. the number DEGs implicated in a particular pathway divided by the total number of DEGs) on the x-axis, and by using split = “.sign” with facet_grid() to split pathways based on enrichment direction. Results (pathway ID, normalized enrichment score, description) from each GSEA KEGG contrast were then concatenated into a data frame and plotted using the pheatmap R package (v1.0.13) [100]. Visualization of selected pathways, such as *ribosomal biogenesis in eukaryotes* in Fig. 3C, from GSEA was achieved using the pathview R package (v1.46.0)[107].

#### Time-Series Soft Clustering Analysis with Mfuzz

Before plotting Mfuzz clusters, variance stabilizing transformation (VST) was applied to the DESeq2 raw count matrix to normalize for sequencing depth. The matrix was then filtered to only include genes with high variation (standard deviation > 0.4) and standardized using the standardise() function within the Mfuzz package (v2.66.0)[41]. The optimal number of clusters was determined using an iterative approach. A range of 4 to 12 clusters was tested, and their corresponding within-cluster error was measured. The number of clusters was chosen based on the point which minimized within-cluster error significantly before reaching a plateau. Enrichment analysis across multiple clusters was then performed using the compareCluster() function and either the GO or KEGG databases[108]. VST-transformed counts were plotted over time for genes relevant to the proposed mechanism.

### Quantitative RT-PCR (qPCR)

Primer pairs for qPCR were designed using Geneious Prime v. 2025.0.3 software (Biomatters, Inc., San Diego, California, USA) for the following genes: peptidylprolyl isomerase A *(PPIA)*, EGF containing fibulin extracellular matrix protein 1 *(EFEMP1)*, niban apoptosis regulator 1 *(FAM129A),* vimentin *(VIM)*, insulin-like growth factor binding protein 5 *(IGFBP5)*, pleckstrin homology-like domain family A member 1 *(PHLDA1)*, claudin 11 *(CLDN11)*, lysophosphatidic acid receptor 1 *(LPAR1)*, isthmin 1 *(ISM1),* WNT1-inducible-signaling pathway protein 2 *(WISP2),* cadherin EGF LAG seven-pass G-type receptor 1 *(CELSR1),* and collagen alpha-1(XII) chain *(COL12A1)*.

Primer pairs were screened for efficiencies between 88 and 112% using a standard curve generated from triplicate 10-fold serial dilutions of cDNA, ranging from 30 ng to 0.003 ng. Diomni™ Design and Analysis v.3.0.1 software (Applied Biosystems, Foster City, California, USA) was utilized to plot the quantification cycle (Cq) against the quantity of DNA and determine the slope of the line. Efficiency was then calculated using the following equation within the software.

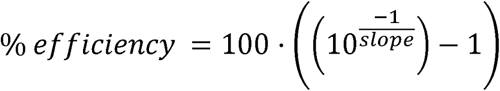

Diomni™ Design and Analysis v.3.0.1 software was also used to visualize melt curves, which confirmed that each primer pair amplified a single, specific amplicon. Successful primer pairs are shown in Supplementary Table 4.

Six independent batches (n = 6) of anchorage-dependent cells were adapted to suspension for qPCR analysis. RNA was isolated and purified from adherent and suspension DF-1s using RNeasy Mini Kit (Qiagen, 74104) and on-column DNase digestion with the RNase-free DNase Set (Qiagen, 79254) according to manufacturer’s instructions. Adherent cells were dissociated from tissue culture plastic using TrypLE to avoid RNA degradation before purification[88].RNA concentration, 260/280 ratios, and 260/230 ratios were determined using NanoDrop™ UV-Vis Spectrophotometer (ThermoFisher, ND-ONE-W). Reverse transcription of 1 ug of total RNA to cDNA was performed with an iScript ™ cDNA Synthesis Kit (Bio-Rad, Hercules, California, USA; 1708890) or High-Capacity RNA-to-cDNA Kit (ThermoFisher; 4387406) according to the manufacturer’s instructions.

After cDNA was synthesized, qPCR was performed on QuantStudio® 5 Real-Time PCR (Applied Biosystems, A28138) with technical triplicates (n = 3) for each biological replicate (n = 6) using the previously described primer pairs and SsoAdvanced Universal SYBR® Green Supermix (Bio-Rad, 1725270) or PowerUp SYBR Green Master Mix for qPCR (ThermoFisher; A25742). Using Diomni™ Design and Analysis v.3.0.1 software, ΔCq was calculated for each sample by subtracting the Cq of the endogenous control gene, *PPIA,* from the Cq of each gene of interest. Next, ΔΔCq was calculated by subtracting the ΔCq of the calibrator sample (i.e. the first adherent biological replicate) from the ΔCq of all other samples. Relative quantification was calculated using the following equation:

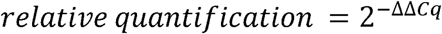

### Statistical Analysis

The data were analyzed using GraphPad Prism version 10.1.1 for Windows (GraphPad Software, La Jolla, CA, USA). The specific analyses include two-tailed unpaired t-test, simple linear regression using ordinary least squares, and calculation of Pearson correlation coefficient using two-tailed significance testing. Error bars represent standard deviations in figures. A p-value of 0.05 was used for statistical significance. All experiments in this study were conducted with a sample size of n ≥ 3.

## Notes

### Competing Interest Statement

The authors have declared no competing interest.

## References

1. Henchion M, Moloney AP, Hyland J, Zimmermann J, McCarthy S. Review: Trends for meat, milk and egg consumption for the next decades and the role played by livestock systems in the global production of proteins. Animal. 2021;15: 100287. doi:10.1016/j.animal.2021.100287

2. Barrow P, Nair V, Baigent S, Atterbury R, Clark M. Poultry Health: A Guide for Professionals. CABI; 2021.

3. Post MJ, Levenberg S, Kaplan DL, Genovese N, Fu J, Bryant CJ, et al. Scientific, sustainability and regulatory challenges of cultured meat. Nat Food. 2020;1: 403–415. doi:10.1038/s43016-020-0112-z

4. Pasitka L, Cohen M, Ehrlich A, Gildor B, Reuveni E, Ayyash M, et al. Spontaneous immortalization of chicken fibroblasts generates stable, high-yield cell lines for serum-free production of cultured meat. Nat Food. 2022; 1–16. doi:10.1038/s43016-022-00658-w

5. Stout AJ, Arnett MJ, Chai K, Guo T, Liao L, Mirliani AB, et al. Immortalized Bovine Satellite Cells for Cultured Meat Applications. ACS Synth Biol. 2023;12: 1567–1573. doi:10.1021/acssynbio.3c00216

6. Stout AJ, Rittenberg ML, Shub M, Saad MK, Mirliani AB, Dolgin J, et al. A Beefy-R culture medium: Replacing albumin with rapeseed protein isolates. Biomaterials. 2023;296: 122092. doi:10.1016/j.biomaterials.2023.122092

7. Radošević K, Dukić B, Andlar M, Slivac I, Gaurina Srček V. Adaptation and cultivation of permanent fish cell line CCO in serum-free medium and influence of protein hydrolysates on growth performance. Cytotechnology. 2016;68: 115–121. doi:10.1007/s10616-014-9760-x

8. Mitić R, Cantoni F, Börlin CS, Post MJ, Jackisch L. A simplified and defined serum-free medium for cultivating fat across species. iScience. 2022;26: 105822. doi:10.1016/j.isci.2022.105822

9. David S, Ianovici I, Guterman Ram G, Shaulov Dvir Y, Lavon N, Levenberg S. Pea Protein-Rich Scaffolds Support 3D Bovine Skeletal Muscle Formation for Cultivated Meat Application. Adv Sustain Syst. n/a: 2300499. doi:10.1002/adsu.202300499

10. Su L, Jing L, Zeng S, Fu C, Huang D. 3D Porous Edible Scaffolds from Rye Secalin for Cell-Based Pork Fat Tissue Culturing. J Agric Food Chem. 2024 [cited 13 May 2024]. doi:10.1021/acs.jafc.3c09713

11. Liu J, DeYoung SM, Zhang M, Zhang M, Cheng A, Saltiel AR. Changes in integrin expression during adipocyte differentiation. Cell Metab. 2005;2: 165–177. doi:10.1016/j.cmet.2005.08.006

12. Adhesion Proteins - An Impact on Skeletal Myoblast Differentiation | PLOS ONE. [cited 26 Nov 2024]. Available: https://journals.plos.org/plosone/article?id=10.1371/journal.pone.0061760

13. Ravikumar M, Powell D. Cell line development and utilisation trends in the cultivated meat industry.

14. Malm M, Saghaleyni R, Lundqvist M, Giudici M, Chotteau V, Field R, et al. Evolution from adherent to suspension: systems biology of HEK293 cell line development. Sci Rep. 2020;10: 18996. doi:10.1038/s41598-020-76137-8

15. Walther CG, Whitfield R, James DC. Importance of Interaction between Integrin and Actin Cytoskeleton in Suspension Adaptation of CHO cells. Appl Biochem Biotechnol. 2016;178: 1286–1302. doi:10.1007/s12010-015-1945-z

16. Paoli P, Giannoni E, Chiarugi P. Anoikis molecular pathways and its role in cancer progression. Biochim Biophys Acta BBA - Mol Cell Res. 2013;1833: 3481–3498. doi:10.1016/j.bbamcr.2013.06.026

17. Dill V, Pfaff F, Zimmer A, Beer M, Eschbaumer M. Adherent and suspension baby hamster kidney cells have a different cytoskeleton and surface receptor repertoire. PLoS ONE. 2021;16: e0246610. doi:10.1371/journal.pone.0246610

18. Huh HD, Sub Y, Oh J, Kim YE, Lee JY, Kim H-R, et al. Reprogramming anchorage dependency by adherent-to-suspension transition promotes metastatic dissemination. Mol Cancer. 2023;22: 63. doi:10.1186/s12943-023-01753-7

19. Dai X, Miao Y, Han P, Zhang X, Yang S, Lv Q, et al. PABPC1 Enables Cells with the Suspension Cultivation Feature. ACS Synth Biol. 2021;10: 309–317. doi:10.1021/acssynbio.0c00440

20. Zhou H, Loo LSW, Ong FYT, Lou X, Wang J, Myint MK, et al. Cost-effective production of meaty aroma from porcine cells for hybrid cultivated meat. Food Chem. 2025; 142946. doi:10.1016/j.foodchem.2025.142946

21. Golkar-Narenji A, Antosik P, Nolin S, Rucinski M, Jopek K, Zok A, et al. Gene Ontology Groups and Signaling Pathways Regulating the Process of Avian Satellite Cell Differentiation. Genes. 2022;13: 242. doi:10.3390/genes13020242

22. Klatt A, Wollschlaeger JO, Albrecht FB, Rühle S, Holzwarth LB, Hrenn H, et al. Dynamically cultured, differentiated bovine adipose-derived stem cell spheroids as building blocks for biofabricating cultured fat. Nat Commun. 2024;15: 9107. doi:10.1038/s41467-024-53486-w

23. Gora K, Goranov A, Katz D, Lamb A-K, Dalvi S, Salinas M, et al. Engineered bovine cell lines for suspension culture. WO2023205677A1, 2023. Available: https://patents.google.com/patent/WO2023205677A1/en

24. Krieger J, Elangovan NB, Ahmed SM. Animal cell line and process development of cultivated meat products. WO2023018995A1, 2023. Available: https://patents.google.com/patent/WO2023018995A1/en?q=(%22cultivated+meat%22+%22bioprocess%22)&oq=%22cultivated+meat%22+and+%22bioprocess%22

25. Wang L, Cai B, Zhou S, Zhu H, Qu L, Wang X, et al. RNA-seq reveals transcriptome changes in goats following myostatin gene knockout. PLOS ONE. 2017;12: e0187966. doi:10.1371/journal.pone.0187966

26. Guo Y, Li S, Na R, Guo L, Huo C, Zhu L, et al. Comparative Transcriptome Analysis of Bovine, Porcine, and Sheep Muscle Using Interpretable Machine Learning Models. Animals. 2024;14: 2947. doi:10.3390/ani14202947

27. Tossolini I, López-Díaz FJ, Kratje R, Prieto CC. Characterization of cellular states of CHO-K1 suspension cell culture through cell cycle and RNA-sequencing profiling. J Biotechnol. 2018;286: 56–67. doi:10.1016/j.jbiotec.2018.09.007

28. Lee N, Shin J, Park JH, Lee GM, Cho S, Cho B-K. Targeted Gene Deletion Using DNA-Free RNA-Guided Cas9 Nuclease Accelerates Adaptation of CHO Cells to Suspension Culture. ACS Synth Biol. 2016;5: 1211–1219. doi:10.1021/acssynbio.5b00249

29. Lee JY, Huh HD, Lee DK, Park SY, Shin JE, Gee HY, et al. Reprogramming anchorage dependency to develop cell lines for recombinant protein expression. Biotechnol J. 2024;19: 2400104. doi:10.1002/biot.202400104

30. UMNSAH/DF-1 - CRL-3586 | ATCC. [cited 1 Nov 2024]. Available: https://www.atcc.org/products/crl-3586

31. Renner WA, Jordan M, Eppenberger HM, Leist C. Cell–cell adhesion and aggregation: Influence on the growth behavior of CHO cells. Biotechnol Bioeng. 1993;41: 188–193. doi:10.1002/bit.260410204

32. O’Neill EN, Cosenza ZA, Baar K, Block DE. Considerations for the development of cost-effective cell culture media for cultivated meat production. Compr Rev Food Sci Food Saf. 2021;20: 686–709. doi:10.1111/1541-4337.12678

33. Ni C, Buszczak M. The homeostatic regulation of ribosome biogenesis. Semin Cell Dev Biol. 2023;136: 13–26. doi:10.1016/j.semcdb.2022.03.043

34. Bendjilali N, MacLeon S, Kalra G, Willis SD, Hossian AKMN, Avery E, et al. Time-Course Analysis of Gene Expression During the Saccharomyces cerevisiae Hypoxic Response. G3 GenesGenomesGenetics. 2017;7: 221–231. doi:10.1534/g3.116.034991

35. Zhang J, Li X, Liu H, Zhou J, Chen J, Du G. Hydrodynamics and mass transfer in spinner flasks: Implications for large scale cultured meat production. Biochem Eng J. 2021;167: 107864. doi:10.1016/j.bej.2020.107864

36. Bodiou V, Cristini N, De Cristofaro L, Pareek T, Rajagopal V, Verrougstraete L, et al. Process intensification of cultivated meat production through microcarrier addition strategy optimisation. Sci Rep. 2025;15: 14080. doi:10.1038/s41598-025-97813-7

37. Cantarero Rivera FJ, Chen J. Computational fluid dynamics modeling of cell cultures in bioreactors and its potential for cultivated meat production—A mini-review. Future Foods. 2022;6: 100195. doi:10.1016/j.fufo.2022.100195

38. Kanehisa M. KEGG: Kyoto Encyclopedia of Genes and Genomes. Nucleic Acids Res. 2000;28: 27–30. doi:10.1093/nar/28.1.27

39. Yue J, López JM. Understanding MAPK Signaling Pathways in Apoptosis. Int J Mol Sci. 2020;21: 2346. doi:10.3390/ijms21072346

40. Mariño G, Niso-Santano M, Baehrecke EH, Kroemer G. Self-consumption: the interplay of autophagy and apoptosis. Nat Rev Mol Cell Biol. 2014;15: 81–94. doi:10.1038/nrm3735

41. Kumar L, E Futschik M. Mfuzz: a software package for soft clustering of microarray data. Bioinformation. 2007;2: 5–7. doi:10.6026/97320630002005

42. Ashburner M, Ball CA, Blake JA, Botstein D, Butler H, Cherry JM, et al. Gene Ontology: tool for the unification of biology. Nat Genet. 2000;25: 25–29. doi:10.1038/75556

43. The Gene Ontology Consortium, Aleksander SA, Balhoff J, Carbon S, Cherry JM, Drabkin HJ, et al. The Gene Ontology knowledgebase in 2023. Baryshnikova A, editor. GENETICS. 2023;224: iyad031. doi:10.1093/genetics/iyad031

44. Alberts B, Johnson A, Lewis J, Raff M, Roberts K, Walter P. Cell Junctions. Molecular Biology of the Cell 4th edition. Garland Science; 2002. Available: https://www.ncbi.nlm.nih.gov/books/NBK26857/

45. Kim D-H, Lee J, Suh Y, Cressman M, Lee SS, Lee K. Adipogenic and Myogenic Potentials of Chicken Embryonic Fibroblasts in vitro: Combination of Fatty Acids and Insulin Induces Adipogenesis. Lipids. 2020;55: 163–171. doi:10.1002/lipd.12220

46. Kim S-H, Kim C-J, Lee E-Y, Hwang Y-H, Joo S-T. Chicken Embryo Fibroblast Viability and Trans-Differentiation Potential for Cultured Meat Production Across Passages. 2024. doi:10.20944/preprints202408.1676.v1

47. Han Z, Ni J, Smits P, Underhill CB, Xie B, Chen Y, et al. Extracellular matrix protein 1 (ECM1) has angiogenic properties and is expressed by breast tumor cells. FASEB J. 2001;15: 988–994. doi:10.1096/fsb2fj990934com

48. Song E, Hou Y, Yu S, Chen S, Huang J, Luo T, et al. EFEMP1 expression promotes angiogenesis and accelerates the growth of cervical cancer *in vivo*. Gynecol Oncol. 2011;121: 174–180. doi:10.1016/j.ygyno.2010.11.004

49. Genovese F, Karsdal MA. Chapter 27 - Type XXVII collagen. In: Karsdal MA, editor. Biochemistry of Collagens, Laminins and Elastin (Third Edition). Academic Press; 2024. pp. 217–221. doi:10.1016/B978-0-443-15617-5.00023-8

50. Zhou J-K, Fan X, Cheng J, Liu W, Peng Y. PDLIM1: Structure, function and implication in cancer. Cell Stress. 5: 119–127. doi:10.15698/cst2021.08.254

51. Liu G, Zhang P, Chen S, Chen Z, Qiu Y, Peng P, et al. FAM129A promotes self-renewal and maintains invasive status via stabilizing the Notch intracellular domain in glioma stem cells. Neuro-Oncol. 2023;25: 1788–1801. doi:10.1093/neuonc/noad079

52. Tang H, Chang H, Dong Y, Guo L, Shi X, Wu Y, et al. Architecture of cell–cell adhesion mediated by sidekicks. Proc Natl Acad Sci. 2018;115: 9246–9251. doi:10.1073/pnas.1801810115

53. Song X, Zhou L, Yang W, Li X, Ma J, Qi K, et al. PHLDA1 is a P53 target gene involved in P53-mediated cell apoptosis. Mol Cell Biochem. 2024;479: 653–664. doi:10.1007/s11010-023-04752-w

54. Kim DJ, Yi YW, Dong Z, Seong Y-S. Therapeutic implication of oxidative stress-induced growth inhibitor 1 (OSGIN1) in cancer. Oncogene. 2025;44: 997–1006. doi:10.1038/s41388-025-03349-5

55. Li C-F, Chen J-Y, Ho Y-H, Hsu W-H, Wu L-C, Lan H-Y, et al. Snail-induced claudin-11 prompts collective migration for tumour progression. Nat Cell Biol. 2019;21: 251–262. doi:10.1038/s41556-018-0268-z

56. Yu S, Wang Y, Yuan X, Yang Y, Wang X, Qin M. A Novel COL12A1 Mutation Causes Oral Tissue Abnormalities by Regulating Gingival Fibroblast Function. Oral Dis. n/a. doi:10.1111/odi.70000

57. Carvalho L, Bergstralh DT, Cristo I, Bosveld F. Editorial: Non-cadherin based cell adhesion in tissue remodeling. Front Cell Dev Biol. 2024;12. doi:10.3389/fcell.2024.1408075

58. Riquelme-Guzmán C, Stout AJ, Kaplan DL, Flack JE. Unlocking the potential of cultivated meat through cell line engineering. iScience. 2024; 110877. doi:10.1016/j.isci.2024.110877

59. Jaluria P, Betenbaugh M, Konstantopoulos K, Shiloach J. Enhancement of cell proliferation in various mammalian cell lines by gene insertion of a cyclin-dependent kinase homolog. BMC Biotechnol. 2007;7: 71. doi:10.1186/1472-6750-7-71

60. Mattson MP. Hormesis defined. Ageing Res Rev. 2008;7: 1–7. doi:10.1016/j.arr.2007.08.007

61. Pinney E, Liu K, Sheeman B, Mansbridge J. Human three-dimensional fibroblast cultures express angiogenic activity. J Cell Physiol. 2000;183: 74–82. doi:10.1002/(SICI)1097-4652(200004)183:1<74::AID-JCP9>3.0.CO;2-G

62. Bellot G, Garcia-Medina R, Gounon P, Chiche J, Roux D, Pouysségur J, et al. Hypoxia-Induced Autophagy Is Mediated through Hypoxia-Inducible Factor Induction of BNIP3 and BNIP3L via Their BH3 Domains. Mol Cell Biol. 2009;29: 2570–2581. doi:10.1128/MCB.00166-09

63. Bristow RG, Hill RP. Hypoxia, DNA repair and genetic instability. Nat Rev Cancer. 2008;8: 180–192. doi:10.1038/nrc2344

64. Su J, Song Y, Zhu Z, Huang X, Fan J, Qiao J, et al. Cell–cell communication: new insights and clinical implications. Signal Transduct Target Ther. 2024;9: 196. doi:10.1038/s41392-024-01888-z

65. Steinbrech DS, Longaker MT, Mehrara BJ, Saadeh PB, Chin GS, Gerrets RP, et al. Fibroblast Response to Hypoxia: The Relationship between Angiogenesis and Matrix Regulation. J Surg Res. 1999;84: 127–133. doi:10.1006/jsre.1999.5627

66. Coppé J-P, Kauser K, Campisi J, Beauséjour CM. Secretion of Vascular Endothelial Growth Factor by Primary Human Fibroblasts at Senescence *. J Biol Chem. 2006;281: 29568–29574. doi:10.1074/jbc.M603307200

67. Boulon S, Westman BJ, Hutten S, Boisvert F-M, Lamond AI. The Nucleolus under Stress. Mol Cell. 2010;40: 216–227. doi:10.1016/j.molcel.2010.09.024

68. Turi Z, Lacey M, Mistrik M, Moudry P. Impaired ribosome biogenesis: mechanisms and relevance to cancer and aging. Aging. 2019;11: 2512–2540. doi:10.18632/aging.101922

69. Montanaro L, Treré D, Derenzini M. Changes in ribosome biogenesis may induce cancer by down-regulating the cell tumor suppressor potential. Biochim Biophys Acta BBA - Rev Cancer. 2012;1825: 101–110. doi:10.1016/j.bbcan.2011.10.006

70. Chadha Y, Khurana A, Schmoller KM. Eukaryotic cell size regulation and its implications for cellular function and dysfunction. Physiol Rev. 2024;104: 1679–1717. doi:10.1152/physrev.00046.2023

71. Vogler M, Vogel S, Krull S, Farhat K, Leisering P, Lutz S, et al. Hypoxia Modulates Fibroblastic Architecture, Adhesion and Migration: A Role for HIF-1α in Cofilin Regulation and Cytoplasmic Actin Distribution. PLOS ONE. 2013;8: e69128. doi:10.1371/journal.pone.0069128

72. Badley RA, Woods A, Carruthers L, Rees DA. Cytoskeleton changes in fibroblast adhesion and detachment. J Cell Sci. 1980;43: 379–390. doi:10.1242/jcs.43.1.379

73. Jimenez-Lopez JC. Cytoskeleton: Structure, Dynamics, Function and Disease. BoD – Books on Demand; 2017.

74. Sjaastad MD, Nelson WJ. Integrin-mediated calcium signaling and regulation of cell adhesion by intracellular calcium. BioEssays. 1997;19: 47–55. doi:10.1002/bies.950190109

75. Clapham DE. Calcium Signaling. Cell. 2007;131: 1047–1058. doi:10.1016/j.cell.2007.11.028

76. Doucet C, Brouty-Boyé D, Pottin-Clemenceau C, Jasmin C, Canonica GW, Azzarone B. IL-4 and IL-13 specifically increase adhesion molecule and inflammatory cytokine expression in human lung fibroblasts. Int Immunol. 1998;10: 1421–1433. doi:10.1093/intimm/10.10.1421

77. Meager A. Cytokine regulation of cellular adhesion molecule expression in inflammation. Cytokine Growth Factor Rev. 1999;10: 27–39. doi:10.1016/S1359-6101(98)00024-0

78. Lin KC, Moroishi T, Meng Z, Jeong H-S, Plouffe SW, Sekido Y, et al. Regulation of Hippo pathway transcription factor TEAD by p38 MAPK-induced cytoplasmic translocation. Nat Cell Biol. 2017;19: 996–1002. doi:10.1038/ncb3581

79. Shao D, Zhai P, Del Re DP, Sciarretta S, Yabuta N, Nojima H, et al. A Functional Interaction between Hippo-YAP Signaling and FoxO1 Mediates the Oxidative Stress Response. Nat Commun. 2014;5: 3315. doi:10.1038/ncomms4315

80. Welsh DJ, Scott PH, Peacock AJ. p38 MAP kinase isoform activity and cell cycle regulators in the proliferative response of pulmonary and systemic artery fibroblasts to acute hypoxia. Pulm Pharmacol Ther. 2006;19: 128–138. doi:10.1016/j.pupt.2005.04.008

81. Mokhtari RB, Ashayeri N, Baghaie L, Sambi M, Satari K, Baluch N, et al. The Hippo Pathway Effectors YAP/TAZ-TEAD Oncoproteins as Emerging Therapeutic Targets in the Tumor Microenvironment. Cancers. 2023;15: 3468. doi:10.3390/cancers15133468

82. Yin Z-T, Zhu F, Lin F-B, Jia T, Wang Z, Sun D-T, et al. Revisiting avian ‘missing’ genes from de novo assembled transcripts. BMC Genomics. 2019;20: 4. doi:10.1186/s12864-018-5407-1

83. Couto PS, Stibbs DJ, Sanchez BC, Khalife R, Panagopoulou TI, Barnes B, et al. Generating suspension-adapted human mesenchymal stromal cells (S-hMSCs) for the scalable manufacture of extracellular vesicles. Cytotherapy. 2024;0. doi:10.1016/j.jcyt.2024.06.011

84. Sène M-A, Xia Y, Kamen AA. Comparative Transcriptomic Analyses of a Vero Cell Line in Suspension versus Adherent Culture Conditions. Int J Cell Biol. 2023;2023: 9364689. doi:10.1155/2023/9364689

85. Kluge S, Benndorf D, Genzel Y, Scharfenberg K, Rapp E, Reichl U. Monitoring changes in proteome during stepwise adaptation of a MDCK cell line from adherence to growth in suspension. Vaccine. 2015;33: 4269–4280. doi:10.1016/j.vaccine.2015.02.077

86. Park JY, Jeong AL, Joo HJ, Han S, Kim S-H, Kim H-Y, et al. Development of suspension cell culture model to mimic circulating tumor cells. Oncotarget. 2017;9: 622–640. doi:10.18632/oncotarget.23079

87. Kumar S, Lazau E, Kim C, N Thadhani N, R Prausnitz M. Serum Protects Cells and Increases Intracellular Delivery of Molecules by Nanoparticle-Mediated Photoporation. Int J Nanomedicine. 2021;16: 3707–3724. doi:10.2147/IJN.S307027

88. Vrtačnik P, Kos Š, Bustin SA, Marc J, Ostanek B. Influence of trypsinization and alternative procedures for cell preparation before RNA extraction on RNA integrity. Anal Biochem. 2014;463: 38–44. doi:10.1016/j.ab.2014.06.017

89. O’Leary NA, Cox E, Holmes JB, Anderson WR, Falk R, Hem V, et al. Exploring and retrieving sequence and metadata for species across the tree of life with NCBI Datasets. Sci Data. 2024;11: 732. doi:10.1038/s41597-024-03571-y

90. Patro R, Duggal G, Love MI, Irizarry RA, Kingsford C. Salmon provides fast and bias-aware quantification of transcript expression. Nat Methods. 2017;14: 417–419. doi:10.1038/nmeth.4197

91. Dobin A, Davis CA, Schlesinger F, Drenkow J, Zaleski C, Jha S, et al. STAR: ultrafast universal RNA-seq aligner. Bioinforma Oxf Engl. 2013;29: 15–21. doi:10.1093/bioinformatics/bts635

92. Andrews S. FastQC: a quality control tool for high throughput sequence data. 2010. Available: https://www.bioinformatics.babraham.ac.uk/projects/fastqc/

93. García-Alcalde F, Okonechnikov K, Carbonell J, Cruz LM, Götz S, Tarazona S, et al. Qualimap: evaluating next-generation sequencing alignment data. Bioinformatics. 2012;28: 2678–2679. doi:10.1093/bioinformatics/bts503

94. Okonechnikov K, Conesa A, García-Alcalde F. Qualimap 2: advanced multi-sample quality control for high-throughput sequencing data. Bioinformatics. 2016;32: 292–294. doi:10.1093/bioinformatics/btv566

95. Srivastava A, Malik L, Sarkar H, Zakeri M, Almodaresi F, Soneson C, et al. Alignment and mapping methodology influence transcript abundance estimation. Genome Biol. 2020;21: 239. doi:10.1186/s13059-020-02151-8

96. Ewels P, Magnusson M, Lundin S, Käller M. MultiQC: summarize analysis results for multiple tools and samples in a single report. Bioinforma Oxf Engl. 2016;32: 3047–3048. doi:10.1093/bioinformatics/btw354

97. R Core Team. R: A language and environment for statistical computing. R Foundation for Statistical Computing, Vienna, Austria.; 2024. Available: https://www.R-project.org/

98. Soneson C, Love MI, Robinson MD. Differential analyses for RNA-seq: transcript-level estimates improve gene-level inferences. F1000Research. 2015;4: 1521. doi:10.12688/f1000research.7563.1

99. Love MI, Huber W, Anders S. Moderated estimation of fold change and dispersion for RNA-seq data with DESeq2. Genome Biol. 2014;15: 550. doi:10.1186/s13059-014-0550-8

100. Kolde R. pheatmap: Pretty Heatmaps. 2010. p. 1.0.13. doi:10.32614/CRAN.package.pheatmap

101. Stephens M. False discovery rates: a new deal. Biostatistics. 2017;18: 275–294. doi:10.1093/biostatistics/kxw041

102. Wickham H. ggplot2: elegant graphics for data analysis. Second edition. Cham: Springer international publishing; 2016.

103. Michał Krassowski, Arts M, Lagger C, Max. krassowski/complex-upset: v1.3.5. Zenodo; 2022. doi:10.5281/ZENODO.7314197

104. Blighe K. EnhancedVolcano. Bioconductor; 2018. doi:10.18129/B9.BIOC.ENHANCEDVOLCANO

105. Yu G, Wang L-G, Han Y, He Q-Y. clusterProfiler: an R Package for Comparing Biological Themes Among Gene Clusters. OMICS J Integr Biol. 2012;16: 284–287. doi:10.1089/omi.2011.0118

106. Wu T, Hu E, Xu S, Chen M, Guo P, Dai Z, et al. clusterProfiler 4.0: A universal enrichment tool for interpreting omics data. Innov Camb Mass. 2021;2: 100141. doi:10.1016/j.xinn.2021.100141

107. Weijun Luo. pathview. Bioconductor; 2017. doi:10.18129/B9.BIOC.PATHVIEW

108. Guangchuang Yu [Aut C. clusterProfiler. Bioconductor; 2017. doi:10.18129/B9.BIOC.CLUSTERPROFILER

