## Supplemental Information for "Comparative Transcriptomics of Adherent and Suspension Chicken Fibroblast Cell Lines for the Optimization of Cultivated Meat Processes"

Supplementary Information

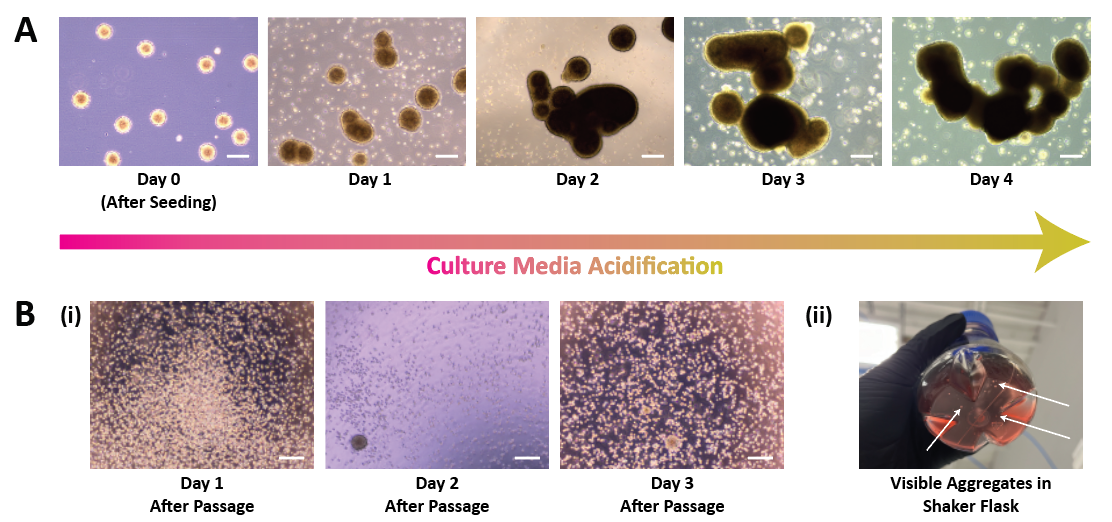

**Supplementary Figure 1:** Morphology of DF-1 cells at T3 and T4 in the suspension adaptation pipeline. (A) Phase contrast images of DF-1 aggregate size increasing over time as spheroids begin to clump together. The color of the culture media in the background of each image turns from pink to yellow over time due to the acidification of phenol red, indicating cell metabolism over time. Scale bars for all images equal 240 μm. (B, i) Phase contrast images of DF-1s in single cell suspension and aggregates during 80 RPM dynamic culture. Scale bars for all images equal 240 μm. (B, ii) Image of cellular aggregates in shaker flask visible to the naked eye.

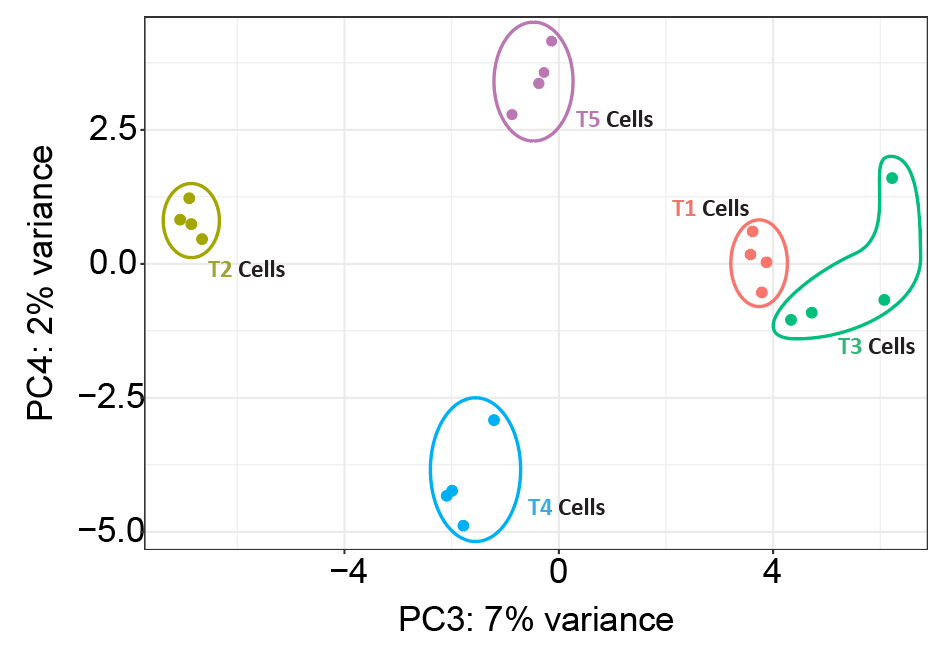

**Supplementary Figure 2:** PCA (PC3 and PC4) of DF-1 cells at each time point in adaptation pipeline; n = 4 for each time point. Pink circles represent adherent T1 cells; olive circles represent T2 spheroids; green circles represent T3 aggregates; blue circles represent T4 cells at 80 RPM; and purple circles represent T5 cells at 100 RPM.

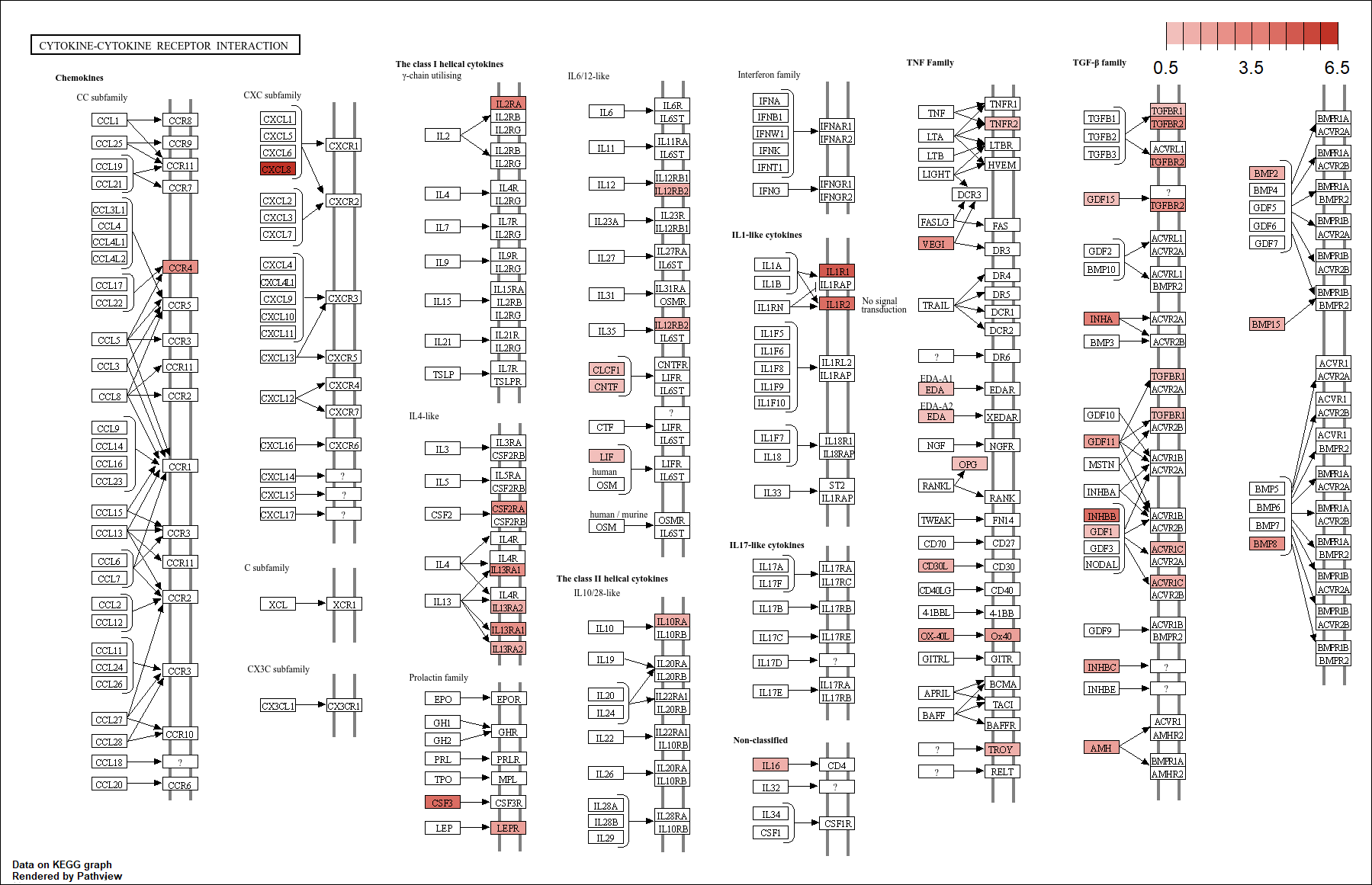

**Supplementary Figure 3:** Pathview image of the cytokine-cytokine receptor interaction pathway, one of the commonly activated pathways across all contrasts against T1. Genes colored in red are upregulated, with darker red colors representing a higher degree of upregulation.

**Supplementary Table 1:** Genes identified from UpSet analysis as being differentially expressed in both the T1 vs. T2 and T1 vs. T3 contrast. Gene symbol, log_2_ fold-change, and adjusted p-value for each contrast is provided.

**Supplementary Table 2:** Genes identified from UpSet analysis as being differentially expressed in all contrasts against adherent baseline (T1 vs. T2 ∩ T1 vs. T3 ∩ T1 vs. T4 ∩ T1 vs. T5). Gene symbol, log_2_ fold-change, and adjusted p-value for each contrast is provided.

**Supplementary Table 3:** Genes assigned to each cluster based on Mfuzz analysis.

| **Gene Name** | **Forward Sequence – 5’ to 3’** | **Reverse Sequence – 5’ to 3’** |
| --- | --- | --- |
| VIM | AGGTGTCACCATGCAGGAAG | ATGTGTCTGCTCCCACACTG |
| EFEMP1 | AGCTATGTTCTTGGCTGCTCT | CCATCTGTGCATTGCGTGTA |
| PHLDA1 | TAGAGCTGCGAGGGGATTCT | GGTGAGCTACCAGCGATACG |
| IGFBP5 | GAGGCTTTATTCTGTCACCACC | ATGCTCTGAGCCACCAACAA |
| CLDN11 | TGACACAGCCACCAAAGAGG | GCGAAACCACCATCATGAGC |
| FAM129A | GGCTGCCAGCTGATCACTTA | GGACTGAGAAACCTGCTCCC |
| LPAR1 | GCCTCCTGAAGACTGTGGTC | CATTGCACTGTGGGCAACAC |
| ISM1 | CAGCTGTCCTTGCTCCTACC | CAGTACCGTGCAGTAGGCTT |
| WISP2 | CAATGAGGAGGGCTGTGAGG | AGAGTGGAACGCAGGTGAAG |
| CELSR1 | AAATCGCTGGACCTGACTGG | GAACTGCCTGTTGTGCACTG |
| COL12A1 | GTGGGCGACAGAACAGTGTA | AGAGGTTTGGTTCTGCCTCG |
| PPIA | TGCCGACGAGAACTTCATCC | TGAAGAACTGGGAGCCGTTC |

**Supplementary Table 4:** List of primer pairs used for qPCR in this study, with their target gene and sequence.

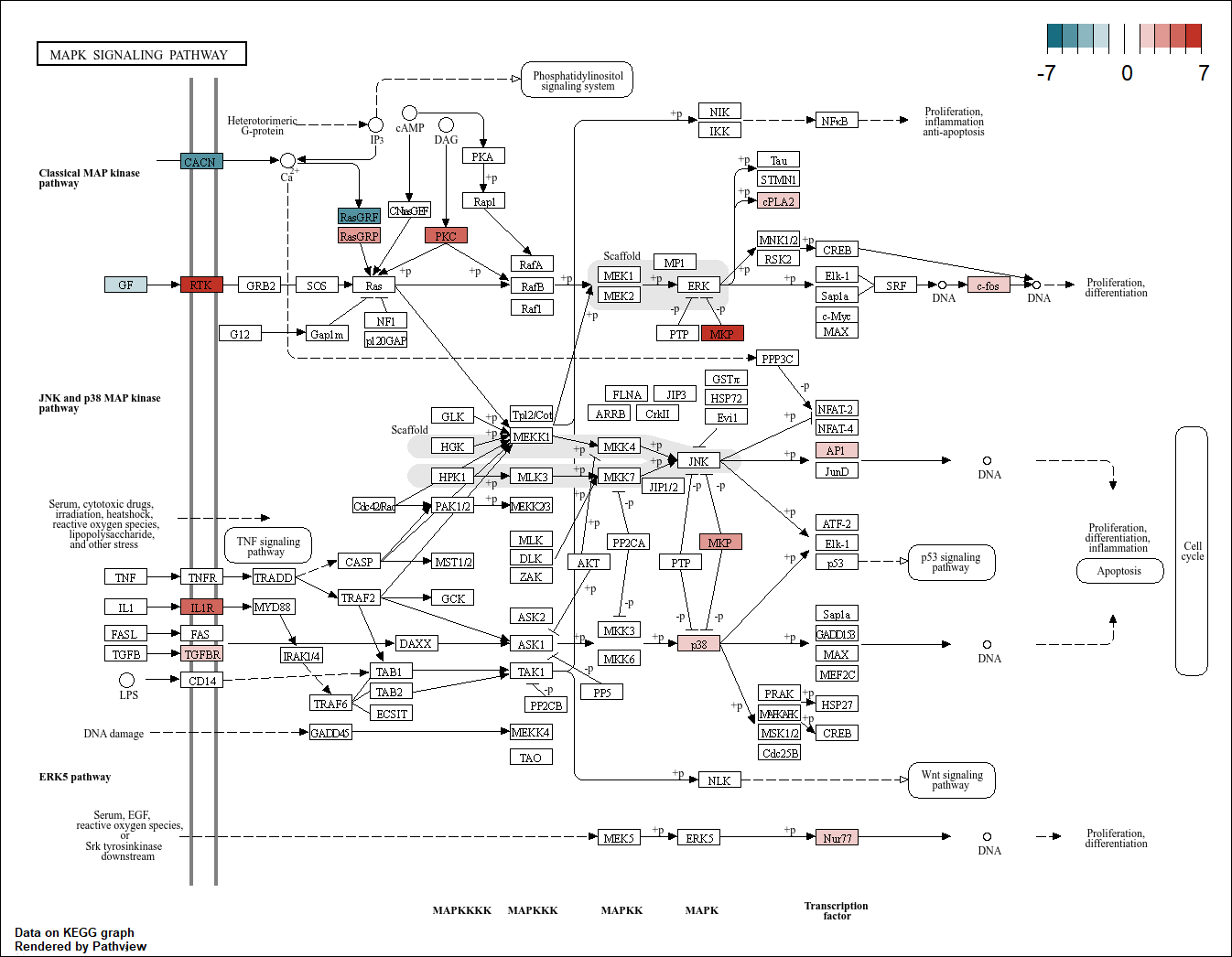

**Supplementary Figure 4:** Pathview image of the MAPK signaling pathway, one of the commonly activated pathways across all contrasts against T1. Genes colored in red are upregulated, with darker red colors representing a higher degree of upregulation. Genes colored in blue are downregulated, with darker blue colors representing a higher degree of downregulation.
